# Single-cell level temporal profiling of tumour-reactive T cells under immune checkpoint blockade

**DOI:** 10.1101/2022.07.19.500582

**Authors:** Jehanne Hassan, Elizabeth Appleton, Bahire Kalfaoglu, Malin Pedersen, José Almeida-Santos, Hisashi Kanemaru, Nobuko Irie, Shane Foo, Omnia Reda, Benjy J.Y. Tan, Il-mi Okazaki, Taku Okazaki, Yorifumi Satou, Kevin Harrington, Alan Melcher, Masahiro Ono

## Abstract

The blockade of the immune checkpoints PD-1 and CTLA-4 enhances T cell response. However, it is largely unknown how antigen-reactive T cells regulate their checkpoint expression *in vivo* and whether and how the checkpoint blockade can change activation dynamics of tumour-reactive T cells. To address this, here we used Nr4a3-*Timer-of-cell-kinetics-and-activity (Tocky)*, which allows analysis of temporal changes of activated T cells following TCR signalling *in vivo*. By analysing melanoma-bearing *Nr4a3* Tocky mice, we elucidate hidden dynamics of tumour-reactive T cells in the steady-state. Checkpoint blockade depleted highly activated effector Treg, while promoting unique effector T cell populations, and thus differentially modulating activation of tumour-reactive T cell populations. Furthermore, multidimensional analysis and seamless analysis of Tocky and scRNA-seq revealed a full spectrum of T cell dynamics in response to tumour burden and treatment with checkpoint blockade. Lastly, we propose a rational design of combinatorial therapy to further enhance T cell activities.

## Introduction

Immune checkpoint blockade (ICB) therapy targeting CTLA-4 and the PD-1/PDL-1 axis has become a mainstay of treatment for many cancers, with the combination particularly successful in melanoma ^1^. CTLA-4 is induced in activated T cells and regulatory T-cells (Treg) and blocks CD28-mediated costimulatory signalling ^2, 3^. PD-1 inhibits proximal TCR signalling molecules and suppresses T cell activation through recruiting the phosphatase SHP2 upon binding to its ligands PD-L1 or PD-L2 on antigen-presenting cells (APCs) and cancer cells ^4–6^. PD-1 expression is induced in T cells by persistent antigenic stimulation, and thus is related to T-cell “exhaustion” in chronic infections and cancer ^7 8^. Blockade of the PD-1 axis by either anti-PD-1 or anti-PD-L1 can inhibit PD-1 binding to PD-L1 and thereby abrogate inhibitory PD-1 signalling ^9^, in addition, the PD-1 signalling pathway may inhibit Treg activities ^10, 11^.

Thus, simultaneous blockade of CTLA-4 and PD-1 has been shown to induce robust anti-tumour T cell responses ^12^, through expanding activated effector CD8^+^ T cells ^13 14^, and a reduction in Treg-mediated immune-suppression ^15^. Currently, however, it is not known what levels of PD-1 or CTLA-4 expression will make T cells vulnerable to ICB-mediated control. Especially, the expression of PD-1 at various levels occurs in a wide range of activated T cells, including early activated T cells, antigen-reactive T cells, exhausted T cells, resident-memory T cells, T follicular helper (Tfh) and T follicular regulatory (T_FR_) cells, and the role of PD-1 may be different between these subsets ^16–20^. In addition, T cells may change their activation and differentiation status in cancer microenvironments ^21, 22^. It is likely that ICB changes the expression levels of immune checkpoint molecules, as CTLA-4 and PD-1 repress genes downstream of T cell activation, including many immune checkpoints ^23^. Furthermore, CTLA-4 and PD-1 can have different functions between different T cell subsets, especially between Treg and non-Treg CD4+ T cells ^24^. These all highlight the importance of single-cell analysis.

Recent advancement of technologies has allowed analysis of single-cell level dynamics of individual T cells, highlighting the importance of understanding temporal aspects of effector and memory T cell differentiation in vivo ^25^. Using scRNA-seq, Wei et al. showed that anti-PD-1 antibody expanded exhausted-like CD8+ T cell subsets, whilst anti-CTLA-4 antibody expanded an ICOS^+^ Th1-like CD4 effector population and exhausted-like CD8^+^ T cells^26^. However, mechanistic understanding of ICB effects on T cells at the single cell level is yet to be fully established ^27^, and will be essential to drive rational use and more effective combinations. Thus, any further significant clinical advances need to be built on dissecting temporal changes of the checkpoint expression at the single-cell level in vivo.

To address these problems, the most straightforward approach is to analyse temporal changes in the expression of checkpoint molecules and other key genes in tumour-reactive T cells at the single-cell level *in vivo*, which should allow identification of tumour-reactive T cell subsets based on their temporally dynamic activities, and elucidate their changes upon checkpoint blockade. <u>T</u>imer-<u>o</u>f-<u>c</u>ell-<u>k</u>inetics-an-activit<u>y</u> (Tocky) uses a fluorescent timer protein which spontaneously and irreversibly changes its emission spectrum from a blue fluorescence (immature) to a red fluorescence (mature) with a maturation half-life of 4 hours ^28^. *Nr4a3* is a TCR downstream gene not expressed in naïve T cells, which is immediately induced specifically by TCR signals ^29 28^. Thus, using *Nr4a3*-Tocky, temporal changes induced by TCR signalling can be revealed by analysing blue and red fluorescence in individual T cells ^28^. Since preceding studies demonstrated that the combination of PD-L1 and CTLA-4 blockade has a therapeutic effect in B16-bearing mice ^30^, we took advantage of this well-established model to reveal hidden dynamics of tumour-reactive T-cells in vivo and their response to ICB.

## MATERIALS AND METHODS

### Animals

*Nr4a3*-Tocky mice, or *BAC Tg (Nr4a3^!1Exon^*^3^*^FTfast^)*, were generated by the Ono group and reported previously ^28^. Briefly, *Nr4a3*-Tocky was generated by a bacterial artificial chromosome (BAC) approach, in which the Fluorescent Timer (FTfast^31^) gene replaced the first coding exon of the *Nr4a3* gene by a knock-in knock-out approach, ensuring that any functional *Nr4a3* protein would not be produced by the transgene. Pronuclear injection was used to generate *Nr4a3-Tocky*. The mice were subsequently bred with *Foxp3*-IRES-GFP (C.Cg-Foxp3tm2Tch/J, Jax ID 006769) and maintained in a C57BL/6 background. All animal experiments were approved by the Animal Welfare and Ethical Review Body at Imperial College London and all animal work was performed in compliance with Home Office and Animal Scientific Procedures Act 1986 in the UK. In addition, all animal experiments in Japan were approved by the Animal Care and Experimentation Committee of the Center for Animal Resources and Development, Kumamoto University, and performed in compliance with Act on Welfare and Management of Animals and Act on the Conservation and Sustainable Use of Biological Diversity through Regulations on the Use of Living Modified Organisms in Japan.

### Tumour inoculation and in vivo antibody administration

The B16-F10 Melanoma cell line was cultured in DMEM 10% FBS 240 mM l-Glutamine and antibiotics (Penicillin and Streptomycin, × 200). 2 × 10^5^ cells were injected subcutaneously in the right flank of Nr4a3 Tocky/wild type C57/Bl6 control mice. Once tumours had reached a palpable size of around 100 mm^3^ (usually day 12 after tumour inoculation), mice were treated intraperitoneally with 0.15 mg anti-CTLA-4 mAb (clone 9H10, BioxCell) or polyclonal Syrian Hamster IgG control (BioxCell) on days 12, 14 and 16 post tumour administration. 0.2mg anti-PD-L1 mAb (clone 1-111A ^32^), or Rat IgG2A Control (clone 2A3, BioxCell) was administered on days 14 and 16, and 0.5 mg OX40 agonist antibody (clone OX-86, BioxCell) or anti Horseradish peroxidase IgG1 Control (clone HRPN, BioxCell), was administered on days 14 and 16. Mice were analysed on day 18 post tumour inoculation.

### Tissue Processing

Tumours were excised using a scalpel, then weighed. These were then placed in an Eppendorf tube and cut into small pieces with dissection scissors. 1ml of digestion buffer composed of 25 μg/mL Liberase (Roche), 250 μl/mL DNASE I (Sigma) 40 μL/mL Trypsin (Sigma) in plain RPMI (Sigma) was added to each tumour, and tubes were then incubated at 37°C on a shaker for 10 minutes, followed by a further 20 minutes at room temperature. The contents of each tube were passed through a 70 μM cell strainer and dissociated using the back of a 5 mL syringe and washed with 5% FBS, 5mM EDTA (Sigma) in plain RPMI for collection in a conical tube. The dissociated tumour cell flow-through was washed twice by adding 10 mL PBS 2 % FBS and centrifugation at 500 × g for 5 minutes. The washed cells were then plated on 96-well plate and stained for flow cytometry.

### Flow Cytometry

Cells were washed by adding 100 μL PBS 2% FBS prior to centrifugation at 800 × g for 2 minutes and the supernatant removed by flicking the plate. The cells were then re-suspended in 50 μL 1:2000 APC-Cy7 Fixable Viability Dye (Invitrogen), or 1:5000 Zombie Aqua Viability dye (BioLegend) and 1:200 Fc block (Invitrogen) in 1×PBS (Gibco) and incubated on ice, in the dark for 15 minutes. Cells were then washed as described previously. In OX40 agonism experiments, the pelleted cells were then re-suspended in 50 μL biotin exclusion cocktail mix including: CD19 (clone ID3), TER-119 (clone TER-119), Gr1 (clone RB6-8C5), CD11b (clone M1/7U), NK1.1 (clone PK136), CD11c (clone N418), TCR-γδ (clone GL3) (all Invitrogen). These were incubated in the dark on ice for 30 minutes, then washed twice. The pelleted cells were then re-suspended in 50 μL of the antibody mix. For single agent and combination experiment, this includes: CD4 BUV395 (GK1.5, BD Biosciences), CD8a BUV737 (53-6.7) from BD Biosciences and TCR-β BV605 (H57-598,), LAG3 BV650 (C9B7W), OX40 BV711 (OX-86), CCR7 BV785 (4B12), CTLA-4 PE (UC10-4B9), PD-1 APC (JK3), CXCR5 PE-Cy7 (SPRCL5), CD44 AF700 (IM7), CD62L (MEL-14), and CD25 PercP-Cy5.5 (PC61) from BioLegend. For OX-40 agonism experiments this includes CD4 BUV395 (GK1.5), CD8a BUV737 (53-6.7), and CD62L BUV805 (MEL-14) from BD Biosciences and TCR-β BV605 (H57-598), ICOS BV650 (398.4A), OX40 BV711 (OX-86), LAG3 BV785 (C9 B7W), KLRG1 PE (2F1), GITR APC (DTA-1), CXCR5 PE-Cy7 (SPRCL5), CD25 AF700 (PC61), Streptavidin PE-Cy5, CD69 PerCP (H1.2F3), and PD-1 APC-Cy7 (29F.1A12) from BioLegend. Antibody cocktails were incubated in the dark on ice for 30 minutes. The samples were then washed twice and re-suspended in 200 μL of PBS 2% FBS, transferred into FACS tubes to be acquired on a Cytek Aurora Flow Cytometer, along with single stains for each antibody used.

### Data Analysis and Statistics

Flow cytometric data analysis was performed using FlowJo version 10 and R version 3.6.3, using the packages *FlowCore* ^33^. Data were visualised using the CRAN packages *ggplot2*, *gplots*, *gridExtra*, *Rmisc,* and *plotrix*. Student’s t-tests, one way and two-way ANOVA was performed using the package *stats*.

### Tocky UMAP analysis

The threshold of Timer fluorescence was set using a fully stained C57/Bl6wild-type T cells, using matched tissues. The algorithms for Forward-Scatter (FSC) normalisation, Tocky Angle transformation, and Tocky Locus analysis were reported previously ^28^. Computational clustering of UMAP data was performed using the CRAN packages *umap* ^34^ and *mclust* ^35^. For UMAP analysis, firstly, angle transformed data (data frame with marker data and Angle and Intensity as added parameters) was generated for each cell type of interest. Marker data used as parameters for UMAP analysis were arc-sinh transformed, and extreme negative outliers were removed. Then UMAP was applied to all concatenated samples, on marker expression only. Importantly, no Timer fluorescence data were included for the UMAP calculation. Heatmaps were generated to uncover marker and timer expression in the UMAP space, using the CRAN package *ggplot2* ^36^. Computational clustering was performed by applying k-means to the PCA result of the variables selected for UMAP. The number of clusters (k number) was determined by Bayesian information criterion (BIC) using *mclust* ^35^.

### Single Cell RNA Sequencing

The scRNA-seq experiment was performed in the facilities in Kumamoto University. Nr4a3-Tocky mice were inoculated with B16-F1 by subcutaneous injection and either anti-PD-L1 (see above) or control IgG (BioXcell BE0089 2A3 rat IgG2a) was administrated on days 0, 3, and 7 (n = 4). On day 8, mice were culled and tdLN cells were pooled for each treatment group and the Tocky populations were flow-sorted using FACS Aria. The three Tocky populations Blue+Red-, Blue+Red+, and Blue-Red+ were sorted in the control group, and the Blue+Red- and Blue-Red+ were sorted from aPD-L1 treated group. The sorted cells were freshly processed with Chromium Single Cell 3’ kit-version 3.1 for library construction. After Tape Station quantification, all the samples were combined to produce the final pool with 6.4 nM. Sequencing was performed using two lanes of HiSeq X Ten at the Macrogen Japan NGS service facility.

### Multidimensional Tocky analysis and ψ-Angle

Briefly, the Tocky quadrant populations (i.e. Blue+Red-, Blue+Red+, and Blue-Red+) are flow sorted and processed for droplet scRNA-seq sequencing, which data are analysed by Principal Component Analysis (PCA) to understand gradual and linear changes of gene expression in individual single cells ^37 38^. A shared nearest neighbour (SNN) modularity optimization-based clustering algorithm is applied to PCA sample scores using the package Seurat ^39^. If successful in the development of Tocky and subsequent sequencing experiments, given that Blue+Red- cells and Blue-Red+ cells have distinct properties, each of them should make a cluster in the PCA space. In PCA analysis, generally, cells with least features are positioned closer to the origin. The more major features cells have, the more distant they are positioned from the origin ^37^. Therefore, using Nr4a3-Tocky, cells distant from the origin of the PCA space will have higher Nr4a3-Timer expression on average. Thus, applying the clustering algorithms, the Seurat cluster of each sample that is the most distant from the origin should show higher Nr4a3 expression, whilst the cluster close to the origin of PCA are mostly Nr4a3 negative and may have mixed Blue+Red- and Blue-Red+ cells. Due to the nature of cells, Blue+Red+ cells should be positioned between the two other Tocky populations. Then, ψ-Angle can be defined as follows.

Firstly, using the barycentre of each of the distant clusters for Blue+Red- and Blue-Red+ cells, ***b*** and ***r***, and that of the origin cluster ***q***, the *New* vector ***n*** and *Arrested* vectors ***a*** are defined in the PCA space:

***n = b - q***

***a = r – q***

Then, the angle 𝜃_*b*_ between 𝙫 and ***α*** is

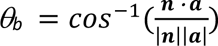

ψ-Angle 𝜃 for any single cells ***x*** between 𝙫 and ***α*** is defined as:

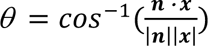

ψ-Angle 𝜃 for any single cells below 0°is defined as 0°, while those above 𝜃*_b_* is defined as 90°. Finally, 𝜃 is further normalised by 𝜃*_b_* and converted into degree, so that 𝜃 has the range 0° - 90°.

For quantification and statistical purposes, ψ-Angle data are categorised into five loci: *New* (Angle 0°), *New-to-Persistent-transitioning* (*NPt*, 0° - 30°), *Persistent* (30° - 60°), *Persistent-to-Arrested-transitioning* (*PAt*, 60° - 90°) and *Arrested* (90°) in the same manner as for Tocky Angle. By definition, ψ-Angle *θ* is ‘unidirectional’ as New vector (Blue+Red-) serves as the origin and Arrested vector (Blue-Red+) as the destination, save for the ‘frequency domain’ Persistent – Arrested, in which Arrested cells can reactivate Nr4a3 transcription upon TCR signalling and ‘move back’ to Persistent ^28^.

## RESULTS

### Tocky reveals hidden dynamics of antigen-reactive T-cells in the steady state of tumour-bearing mice

Our major hypothesis in this study was that effects of the checkpoint blockade on individual tumour-reactive T cells are dependent on the following two factors: (1) whether and how frequently T cells engage with cognate antigen in vivo; and (2) their expression of checkpoint molecules, which can be dynamically altered by TCR signalling. To this end, we investigated T cells from B16-bearing Nr4a3-Tocky mice.

*Nr4a3* is a gene downstream of cognate antigen signals ^28^. Nr4a3-Tocky carries the bacterial artificial chromosome (BAC) *Nr4a3^Timer^* transgene, with its main exon replaced with a Fluorescent Timer gene, and can reveal temporal changes following TCR signals in T cells. Using Nr4a3-Tocky, antigen-reactive T cells, which have recognised their cognate antigen, will express Timer proteins ^28^. (**Fig. 1a**). To understand temporal changes of individual T cells upon receiving TCR signalling, the maturation of Timer fluorescence is analysed by a trigonometric transformation, which produces two variables: Timer Angle and Timer Intensity (**Fig. 1b**). Timer Angle is the angle from the blue axis and Timer Intensity is the distance from Timer-negative cells. For quantification and statistical purposes, Angle data are categorised into five loci: *New* (Angle 0°), *New-to-Persistent-transitioning* (*NPt*, 0° −30°), *Persistent* (30° - 60°), *Persistent-to-Arrested-transitioning* (*PAt*, 60° - 90°) and *Arrested* (90°) as reported before ^28^.

**Figure 1.**
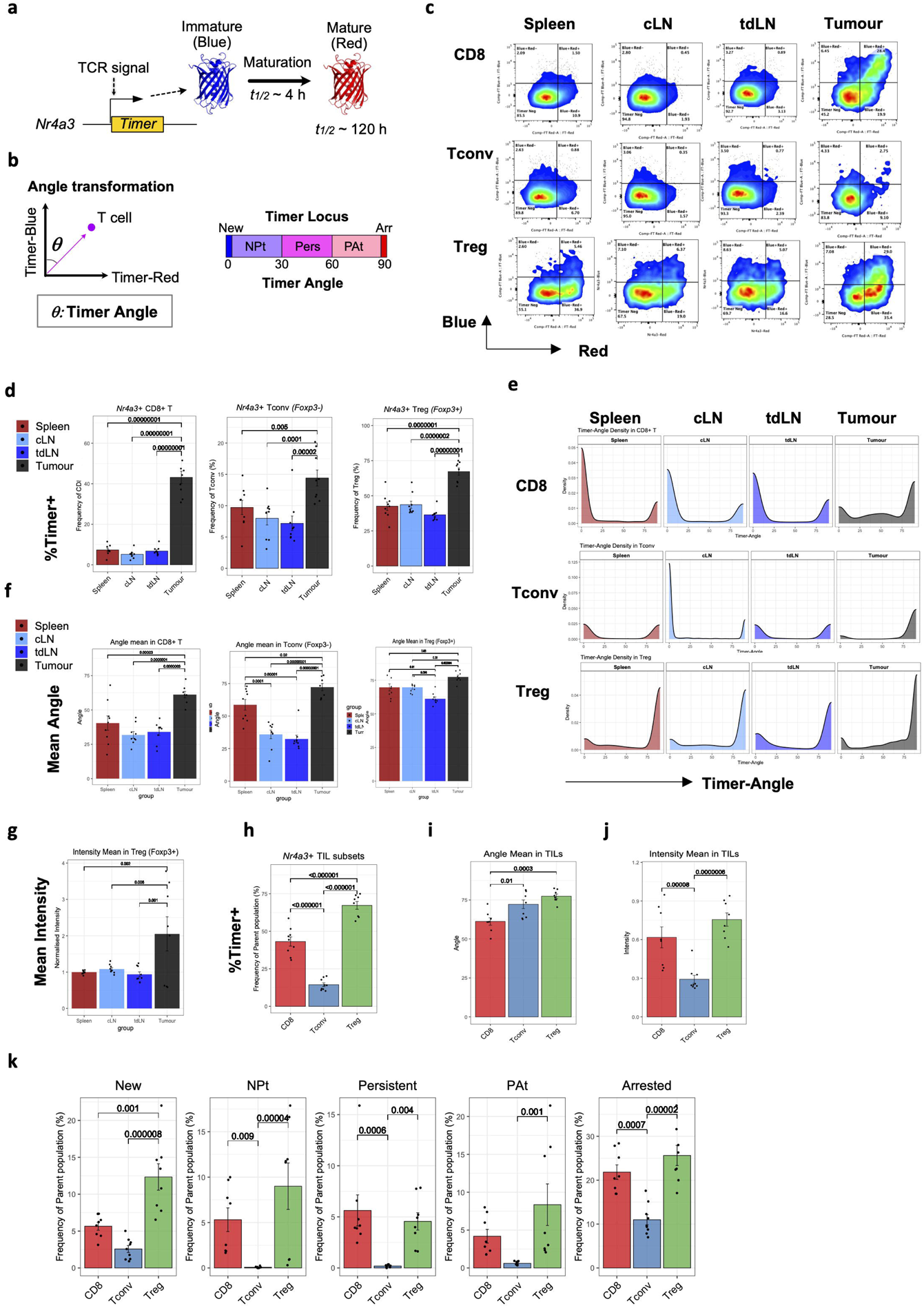
Nr4a3 Tocky reveals hidden dynamics of antigen-reactive T-cells in the steady state of tumour-bearing mice. (**a**) The principle of Nr4a3-Tocky is depicted. TCR signals activate the transcription of *Nr4a3 ^Timer^*, which is translated into immature Timer proteins with a blue fluorescence (Blue). The immature chromophore spontaneously and irreversibly mature into a mature chromophore, which emits a red fluorescence (Red), which maturation half-life is 4 hours. The mature protein is stable with the half-life > 120 hours. (**b**) Rationale for the analysis of Timer Fluorescence data using Angle transformation and Tocky locus categorisation. Trigonometric transformation produces Timer Angle and Intensity. Timer Angle is categorised into the following five loci: *New* (Angle 0°), *New-to-Persistent-transitioning* (*NPt*, 0° - 30°), *Persistent* (Pers, 30° - 60°), *Persistent-to-Arrested-transitioning* (*PAt*, 60° - 90°) and *Arrested* (Arr, 90°). (**c**) Representative flow cytometry plots showing Timer Blue and Red fluorescence in Spleen, tumour draining lymph Node (tdLN), contra-lateral lymph node (cLN) and tumour-infiltrating T cells. (**d**) Bar plots showing the percentage of Nr4a3 Timer+ cells in CD8+ T cells (left), Tconv (middle) and Treg (right) in Spleen, cLN, tdLN and Tumour. (**e**) Density plots (smoothened histograms) showing the abundance of cells in each Angle value. CD8+ T cells, Foxp3- CD4+ T cells (conventional T cells, Tconv) and Treg from the spleen, contra-lateral lymph node (cLN), tumour-draining lymph Node (tdLN) and Tumour are shown. Representative data of three independent experiments (n=3 each). (**f**) Bar plots showing the mean Angle of Nr4a3 Timer+ cells in CD8+ T cells (left), Tconv (middle) and Treg (right) in Spleen, cLN, tdLN and Tumour. (**g**) Bar plots showing the mean Intensity of Nr4a3 Timer+ cells in Treg in Spleen, cLN, tdLN and Tumour. (**h-j**) Bar plots showing (h) the percentage, (i) mean Angle, or (j) mean Intensity of Nr4a3 Timer+ cells from tumour-infiltrating Treg. (**k**) Bar plots showing the percentage of tumour-infiltrating CD8+ T cells, Tconv and Treg in each Tocky Locus out of the parent T cell population. Data were pooled from 3 independent experiments (n = 9 per group in total). Two-way ANOVA with Tukey’s multiple comparisons test was applied. Dots represent individual data points, error bars represent +/- standard error.

By analysing B16-bearing Nr4a3-Tocky mice, CD8^+^ T cells, Foxp3^+^ CD4^+^ T cells (Treg) and Foxp3^-^ CD4^+^ T cells (Tconv) showed unique Timer expression profiles in the spleen, tumour-draining and contra-lateral lymph nodes (tdLN and cLN, respectively), and tumour-infiltrating lymphocytes (TIL) (**Fig. 1c**). Notably, Timer expression occurred in ∼8% of CD8+ T cells, ∼10% of Tconv, and ∼40% of Treg from the spleen and cLN, which is comparable with our previous finding that showed equivalent Timer expression in Treg and naturally arising memory T cells in untreated, healthy mice, indicating that the majority of these T cells react to self-antigens or microbiome antigens ^28^. As expected, tumour-infiltrating T cells had the highest frequency of antigen-reactive T cells as Timer^+^ cells (**Fig. 1c, 1d**).

Next, the flow cytometric Timer fluorescence data was further analysed by Angle transformation (**Fig. 1e**). Tumour-infiltrating T cells were the most enriched with antigen-experienced T cells, showing the highest mean values of Timer Angle (**Fig. 1f**). Notably, T cells in the PAt and Arrested Loci, which were antigen-experienced but not engaging with antigen in a real-time manner as were T cells in the Persistent locus, were remarkably increased in all three populations of tumour-infiltrating T cells. T cells at the Persistent locus, which frequently recognised antigen, were also increased in Treg and CD8^+^ T cells (∼6% in tumour-infiltrating CD8+ T cells, vs ∼0.1% in CD8+ T cells in the LNs and the spleen; ∼5% in tumour-infiltrating Treg vs 1 – 2.5% in Treg in the LNs and the spleen), while Tconv did not show this feature (**Extended Data 1a – 1c**). These data indicate that antigen-experienced T cells (including CD8^+^ T cells, Tconv, and Treg) were accumulated in the tumour, yet the majority only infrequently recognised their cognate antigen.

Notably, tumour-infiltrating Treg had remarkably high Timer Intensity, indicating that they received ‘strong’ or frequent TCR signals, which accumulated Timer protein (**Fig. 1g**). Since the majority of Treg in tumour only infrequently engage with antigen (Extended Data 1c), this result suggests that, although only a small fraction of these Treg were actively engaged with antigen at the time of analysis, the majority of tumour-infiltrating Treg had experienced strong and relatively short antigen engagement and thereby accumulated Timer proteins (note that the half-life of Red is ∼5 days ^40^). In fact, the majority of tumour-infiltrating Treg expressed Timer, while nearly a half of CD8^+^ T cells and only ∼13% of Tconv expressed Timer (**Fig. 1h**). Tconv and Treg had higher Angle than CD8^+^ T cells in the Tumour, indicating that CD8^+^ T cells more frequently recognised antigen, on average, than the CD4+ T cell populations (**Fig. 1i**). Tconv had lower intensity mean than Treg and CD8+ T cells in the Tumour, indicating that Tconv received ‘weaker’ or less frequent TCR signals on average (**Figure 1j**). Interestingly, Tconv had the lowest percentages of Timer+ cells across all the Tocky loci, indicating that Treg and CD8+ T cells dominated antigen recognition events within tumour. The percentage of cells at the New locus was the highest in tumour-infiltrating Treg in comparison to CD8+ T cells and Tconv (**Fig. 1k**).

Collectively, Tocky revealed that a significant proportion of T cells in tumour-bearing mice had actively recognised their cognate antigen and were signalling through their TCR at the time of analysis. Intratumoural T cells were the most enriched with antigen-reactive T cells, and Treg appear to dominate access to antigen within the tumour microenvironment, with the majority receiving TCR signals compared to a lower proportion of CD8 and FoxP3-Tconv.

### Identification of tumour-reactive T-cell populations by *Nr4a3*-Tocky

Next, to identify and reveal marker features of tumour-reactive T cell populations, we applied Uniform Manifold Approximation and Projection (UMAP) and cross-analysed data from cLN and tdLN from non-ICB treated mice, aiming to identify the single cell Treg, Tconv and CD8+ T populations that were increased by the tumour burden (**Fig. 2**).

**Figure 2.**
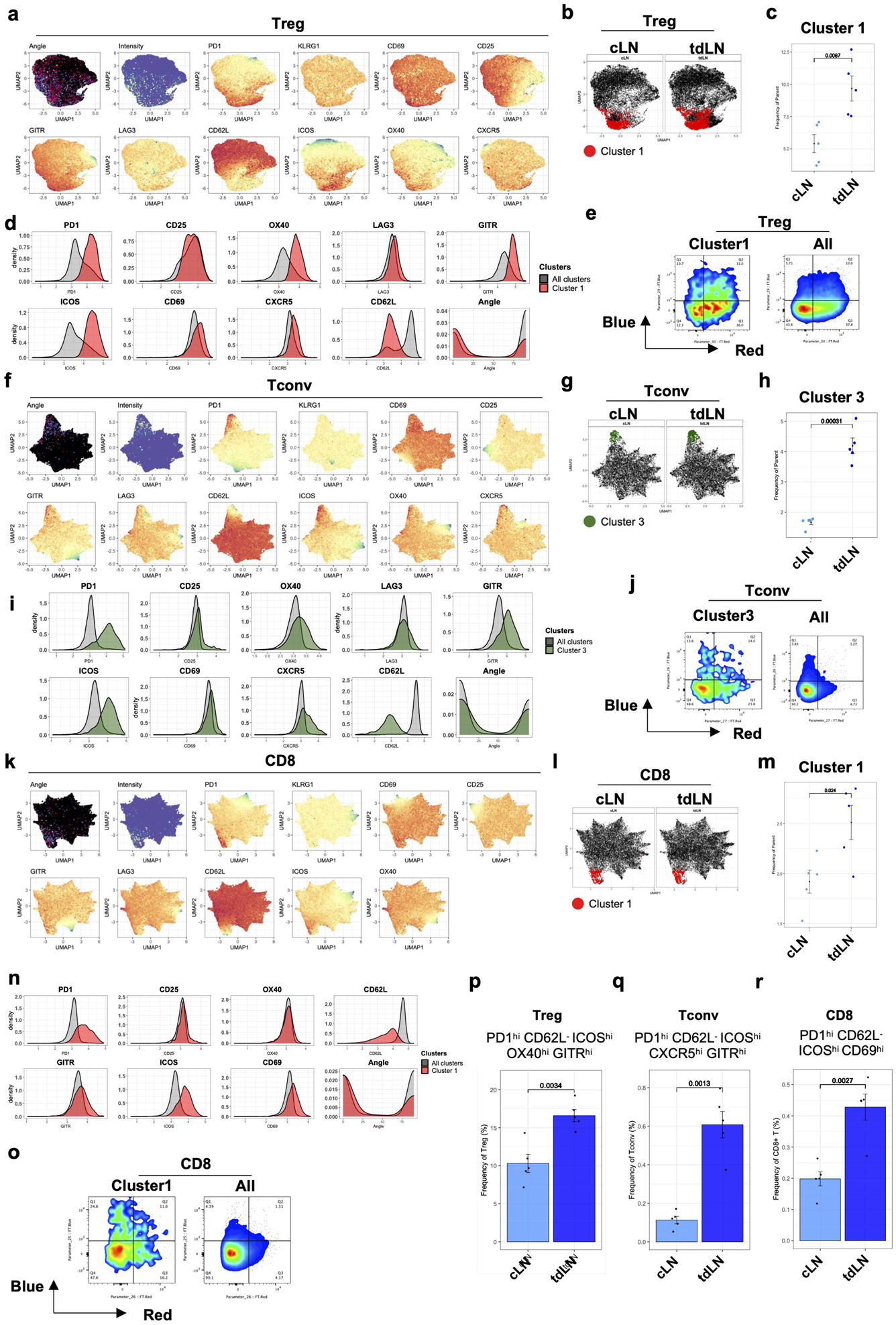
Identification of tumour-reactive T-cell populations by *Nr4a3*-Tocky. UMAP analysis was applied to the data in Fig. 1. (**a**) Heatmaps showing the UMAP analysis of Treg cells from tumour bearing Nr4a3-Tocky mice. The heat colours show Angle, Intensity, and the expression of indicated markers. Briefly, UMAP was applied to Treg cell data from both tumour-draining and contra-lateral lymph nodes (tdLN and cLN, respectively) in B16-F10 bearing mice (n = 5 each). UMAP was performed using KLRG1, CD69, CD25, GITR, LAG3, CD62L, ICOS, OX40 and CXCR5 as input data. (**b**) Cluster 1 cells are highlighted, showing tdLN and cLN cells. (**c**) Scatter plot showing the percentage of Treg in Cluster 1 from either tdLN or cLN (n = 5). (**d**) Overlaid histograms showing the indicated marker expression in Cluster 1 and all concatenated Clusters.(**e**) Timer-Blue and -Red of Treg cells in Cluster 1 and all concatenated clusters.(**f**) Heatmaps showing the UMAP analysis of Tconv. The heat colours show Angle, Intensity, and the expression of indicated markers in Tconv cells from pooled tdLN and cLN cells (n = 5). UMAP was performed using KLRG1, CD69, CD25, GITR, LAG3, CD62L, ICOS, OX40 and CXCR5 as input data. (**g**) Cells in Cluster 3 are highlighted by green, showing tdLN and cLN cells. (**h**) Scatter plot showing the percentage of Treg in Cluster 3 from either tdLN or cLN (n = 5). (**i**) Overlaid histograms showing the expression of indicated markers in Cluster 3 and all concatenated clusters. (**j**) Timer-Blue and -Red of Tconv cells in Cluster 3 and all concatenated clusters. (**k**) Heatmaps showing the UMAP analysis of CD8+ T cells. The heat colours show Angle, Intensity, and the expression of indicated markers in Tconv cells from pooled tdLN and cLN cells. UMAP was performed using KLRG1, CD69, CD25, GITR, LAG3, CD62L, ICOS and OX40 as input data. (**l**) Cells in Cluster 1 are highlighted by red, showing tdLN and cLN. (**m**) Scatter plot showing the percentage of CD8+ T cells in the UMAP Cluster 1 from either tdLN or cLN (n = 6). (**n**) Overlaid histograms showing the expression of indicated markers in Cluster 3 and all concatenated clusters. (**o**) Timer-Blue and -Red of CD8+ T cells in Cluster 3 and all concatenated clusters. (**p**) Bar plot showing the frequency of PD1^+^ CD62L^-^ ICOS^hi^ CXCR5^hi^ GITR^hi^ Tconv in cLN and tdLN. (**q**) Bar plot showing the frequency of PD1^+^ CD62L^-^ ICOS^hi^ GITR^hi^ OX40^hi^ Treg in cLN and tdLN. (**r**) Bar plot showing the frequency of PD1^+^ CD62L^-^ ICOS^hi^ CD69^hi^ CD8 T cells in cLN and tdLN. All data are from the cLN or tdLN of B16-F10 tumour bearing mice (n = 5). Student’s t-test was employed. Dots represent individual data points. Error bars represent +/- Standard error.

First, we aimed to identify the Treg single cell clusters that were increased in tdLN in comparison to cLN. Heatmap analysis of UMAP showed that the expression of PD-1, CD25, GITR, CD69 and OX40 was correlated with Angle and Intensity (**Fig. 2a**). A computational clustering analysis identified 9 clusters (**Extended Data 2a**). Among the 9 clusters, Cluster 1 only showed a significantly increase in tdLN (**Fig. 2b, 2c**). Treg cells in Cluster 1 were characteristically PD-1^hi^ CD25^int^ OX40^hi^ LAG3^+^ GITR^hi^ ICOS^hi^ CD69^+^ CXCR5^hi^ CD62L^lo^; this profile was partially similar to those of T follicular regulatory (T_FR_) cells (i.e. PD-1^hi^ ICOS^hi^ CXCR5^hi^ Treg) and effector Treg (eTreg, ICOS^hi^ GITR^hi^ OX40^hi^ CD62L^lo^ Treg), which are activated Treg with enhanced suppressive activities ^41^ (**Fig. 2d**). Intriguingly, Cluster 1 cells were enriched with antigen-reactive T cells, as the percentage of Timer^+^ cells was 78% and 57% in Cluster1 and all concatenated clusters, respectively (**Fig. 2e**).

Similarly, UMAP analysis showed that Tconv cells with high PD-1, GITR, ICOS, OX40 and CXCR5 expression had higher Angle and Intensity (**Fig. 2f**). Computational clustering identified 9 clusters, and Cluster 3 only was significantly increased in tdLN (**Extended Data 2b** and **Fig. 2g, 2h**). Tconv cells in Cluster 3 were uniquely PD-1^hi^ GITR^hi^ ICOS^hi^ CXCR5^hi^ LAG3^int^ CD25^int^ OX40^int^ CD69^hi^ (**Fig. 2i**), and showed high levels of antigen reactivity (50% in Cluster 3 vs 10% in all other concatenated clusters. **Fig. 2j**).

The analysis of CD8 T cells identified 9 clusters, among which Cluster 1 only was increased in tdLN compared to cLN (**Extended Data 2c** and **Fig. 2l, 2m**). CD8 T cells in Cluster 1 were characteristically PD-1^hi^ GITR^+^ ICOS^hi^ CD69^hi^ CD62L^lo^ with high antigen reactivity (**Fig. 2n, 2o**).

Next, we aimed to validate and identify the significantly increased UMAP clusters using conventional gating approaches. Consistent with the UMAP analysis, we successfully identified the novel T cell populations that were increased upon tumour burden, including the following populations increased in the tdLN compared to the cLN : the Treg populations PD-1^+^ CD62L^-^ and PD-1^+^ CD62L^-^ ICOS^hi^ GITR^hi^ OX40^hi^ (**Extended Data 2e, Fig. 2p**), the Tconv populations PD-1^+^ CD62L^-^ and PD-1^+^ CD62L^-^ ICOS^hi^ CXCR5^hi^ GITR^hi^ (**Extended Data 2d, Fig. 2q**), and the CD8+ T cell populations PD-1^+^ CD62L^-^ and PD-1^+^ CD62L^-^ ICOS^hi^ CD69^hi^ (**Extended Data 2f, Fig. 2r**). Interestingly, in all three T cell subtypes (Treg, Tconv and CD8), PD-1^+^ CD62L^-^ T cells were increased in tdLN relative to cLN (**Extended Data 2g-S2i**)

Collectively, the tumour burden increased antigen-reactive Nr4a3-Timer^+^ PD-1^+^ CD62L^-^ T cells, especially their subfractions ICOS^hi^ GITR^hi^ OX40^hi^ T_FR_-like eTreg population, the ICOS^hi^ CXCR5^hi^ GITR^hi^ T follicular helper (T_FH_)-like cells, and ICOS^hi^ CD69^hi^ CD8+ T cells. Overall, PD-1, ICOS and CD69 expression positively correlated with Timer intensity in CD8^+^ T cells (**Fig. 2k**, **Extended Data 2j**).

### TCR signalling dynamically controls the expression of immune checkpoint and costimulatory molecules on tumour-infiltrating T cells in vivo

Next, we tested if immune checkpoint expression on tumour-infiltrating T cells is dependent on how T cells have recently recognized cognate antigen and received TCR signalling *in vivo*.

PD-1 expression in tumour-infiltrating Treg was increased upon Persistent antigen signalling and was sustained towards the Arrested locus, when Treg rarely recognise antigen (**Extended Data 3a**). In contrast, the expression of CD25, OX40, GITR, ICOS and LAG3 was increased upon persistent antigenic stimulation but decreased in the Arrested locus, indicating that their expression is dependent on real-time activities of TCR signalling (**Extended Data 3a**). As Timer Intensity measures the accumulation of Timer proteins, Timer Intensity in Nr4a3-Tocky shows the history of TCR signalling over time. In tumour-infiltrating Treg, Tocky Intensity was correlated with the expression of PD-1, CD25, OX40, GITR, ICOS and LAG3 (**Extended Data 3b**). In tumour-infiltrating Tconv, the expression of PD-1, OX40 LAG3 and ICOS was increased following Persistent antigenic stimulation, decreased in the Arrested locus (**Extended Data 3c**) and positively correlated with Timer Intensity (**Extended Data 3d**).

In CD8 T cells, the expression of PD-1, LAG3, ICOS, GITR, CD25 and OX40 was increased following persistent antigenic signalling, then decreased in ‘Arrested’ cells (**Extended Data 3e**). Timer Intensity in Tumour-infiltrating CD8+ T cells showed positive correlations with the expression of PD-1, LAG3, ICOS, GITR, CD25 and OX40, whilst it had a negative correlation with CD62L (**Extended Data 3f**).

Collectively, the data above show that tumour infiltrating T cell change their phenotype according to how frequently they have recognised antigen *in vivo*. The expression of key surface markers for T cells, including CD25, OX40, LAG3, ICOS and GITR, was dependent on active TCR signalling. PD-1 expression was sustained only when Tconv and CD8+ T cells persistently receive TCR signalling, while PD-1 expression was uniquely sustained in Treg, not only during active TCR signalling, but also after their detachment from antigen. Collectively, the expression of immune checkpoints, costimulatory molecules, and Treg-associated molecules are dynamically controlled by TCR signalling-driven T cell activation.

### Single cell RNA-seq reveals the full spectrum of gene regulation in tumour-reactive T cells according to TCR signalling

The results above collectively suggest that individual T cells upregulate the expression of activation markers such as PD-1, ICOS, and CD69 as they recognize cognate antigen *in situ*. This leads to the hypothesis that each of the tumour-reactive T cell populations is in a unique status of activation following antigen recognition, dynamically changing their phenotype even in the steady-state tumour. To address this, we performed single cell (sc) RNA-seq to analyse the full spectrum of gene regulation and fully characterize the activation status of tumour-reactive T cells, focusing on the tdLN of B16-bearing Nr4a3-Tocky mice, since tumour-reactive T cells were successfully identified and characterised in tdLN (Fig. 2), where antigen recognition leads to CD8 priming.

Although droplet library construction (e.g. Chromium) allows analysis of tens of thousands of single cells, it cannot obtain Timer fluorescence data for individual single cells. Thus, we developed a new approach, *multidimensional Tocky analysis using* ψ*-Angle* for scRNA-seq of Tocky T cells, in order to seamlessly analyse and integrate Tocky profiles and scRNA transcriptomes (**Fig. 3a**). Briefly, the Tocky quadrant populations (i.e. Blue+Red-, Blue+Red+, and Blue-Red+) are flow sorted and processed for droplet scRNA-seq sequencing, which data are analysed by Principal Component Analysis (PCA) for dimensional reduction. Single-cell level heterogeneity allows bridging of ‘gaps’ between the three distinct Tocky populations, identifying the *New* and *Arrested* vectors in the PCA ^42^. Finally, trigonometric transformation is applied to the PCA data to obtain ψ-Angle, which is the angle from the New vector normalised so that ψ-Angle of Arrested vector is 90° (**Fig. 3a**). Thus, single-cell level heterogeneity and variations in the PCA space allows ‘gaps’ between the three distinct Tocky populations to be bridged. Finally, trigonometric transformation is applied to the PCA data to obtain λφλ-Angle, which is the angle from the New vector normalised so that ψ-Angle will have values between 0° and 90° (**Materials and Methods**)

**Figure 3:**
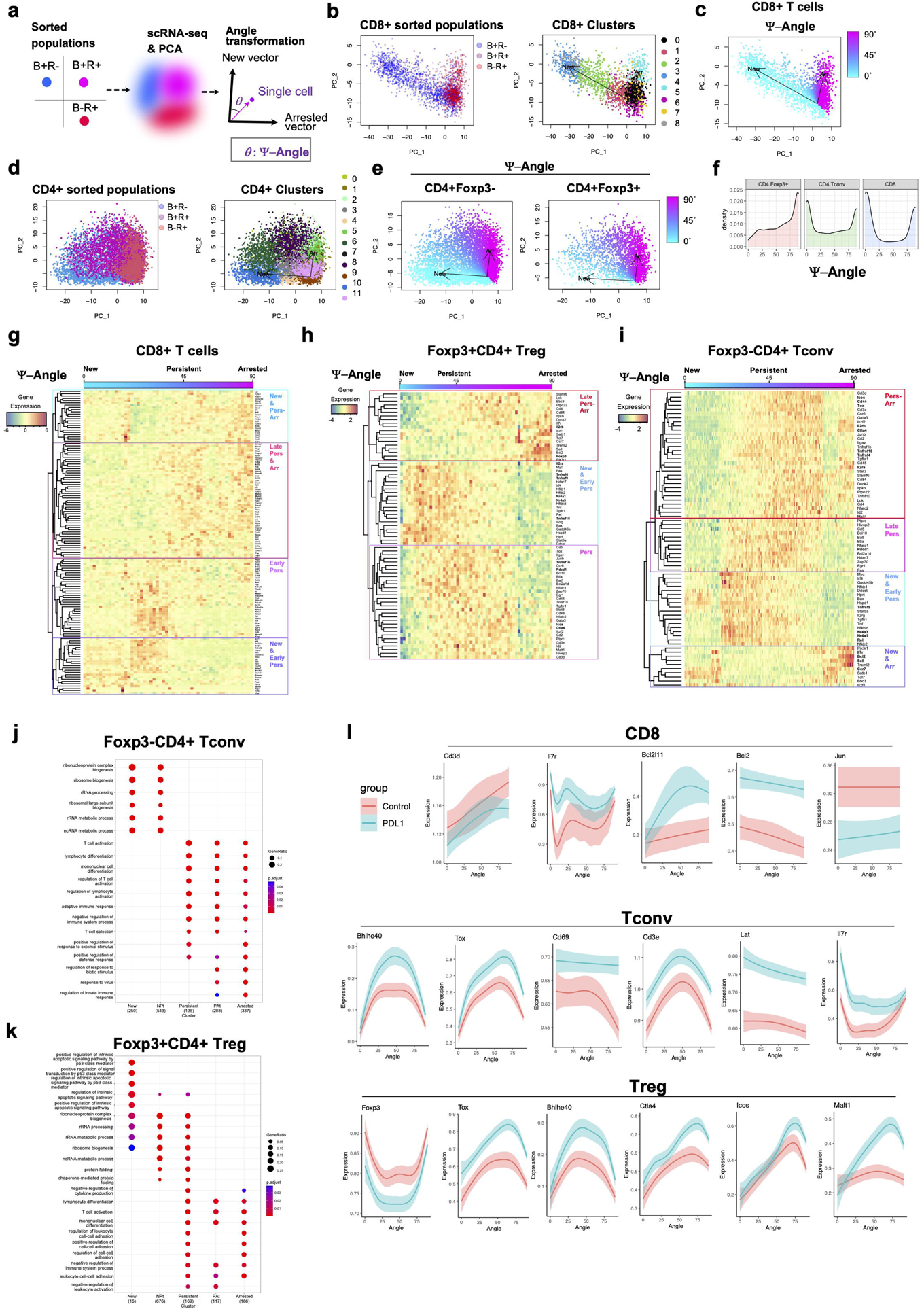
Multidimensional Tocky-single cell RNA-seq analysis reveals the effects of anti-PD-L1 antibody on in vivo dynamics of tumour-reactive T cells. Blue+Red- (B+R-), Blue+Red+ (B+R+), and Blue-Red+ (B-R+) TCRβ+ viable single cell T cells were sorted from B16-F10-bearing Nr4a3-Tocky mice. Samples were pooled for a droplet single cell RNA-seq. The major T cell populations were in silico sorted and further analysed for reconstituting in vivo Tocky dynamics as ψ-Angle. **(a)** Rationale for reconstituting the full spectrum of in vivo dynamics of Tocky maturation using the three flow-sorted populations and Principal Component Analysis (PCA) together with ψ-Angle transformation. **(b)** CD8+ single cells flow-sorted and in silico purified, showing the original sample identities (left) and computational clustering (right). **(c)** ψ-Angle transformation of CD8 T cell data visualised by a heatmap. **(d)** CD4+ single cells flow-sorted and in silico purified, showing the original sample identities (left) and computational clustering (right). **(e)** ψ-Angle transformation of CD4 T cell data visualised by a heatmap for Foxp3- (Tconv, left) and Foxp3+ (Treg, right). **(f)** Density plot of ψ-Angle in CD4 Foxp3+ (Treg), CD4 Foxp3- (Tconv) and CD8 T cells. (**g – i**) Heatmap of gene expression of selected genes in (**g**) CD8+ T cells, (**h**) Treg, and (**i**) Tconv. X axis is ordered by ψ-Angle, showing the average expression of each 40 single cells for each gene. (**j – k**) Pathway analysis for (**j**) Tconv and (**k**) Treg in each ψ-Tocky Locus. (**l**) The expression of DEGs between the aPD-L1 treated and control groups are shown by line graphs with a confidence interval using a Loess smoothing.

PCA of sorted CD8+ T cell Tocky populations showed distinct Blue+Red- and Blue-Red+ clusters, while Blue+Red+ cells positioned in the middle of the two populations, identifying clusters in the multidimensional space (**Fig. 3b**). Accordingly, the ψ-Angle approach can be applied to the data (**Fig. 3c**). Similarly, PCA of sorted CD4+ T cell Tocky populations showed distinct Blue+Red- and Blue-Red+ clusters, with Blue+Red+ cells again positioned in the middle, identifying clusters in the multidimensional space (**Fig. 3d**). Subsequently, ψ-Angle was calculated for these CD4+ T cells, including Foxp3+ and Foxp3- cells (**Fig. 3e**). Interestingly, Foxp3+CD4+ Treg have higher ψ-Angle values than Foxp3-CD4+ Tconv and CD8+ T cells (**Fig. 3f**), which is consistent with the result from flow cytometric Tocky analysis (**Fig. 1e**).

Heatmap analysis of gene expression across ψ-Angle showed that sets of genes were progressively activated and repressed as Timer proteins matured and ψ-Angle increased (**Fig. 3g – 3i**), supporting dynamic gene regulation in T cells upon antigen recognition in the steady-state. Similarly, temporal changes of gene expression profiles in Foxp3^-^CD4^+^ Tconv and Foxp3^+^CD4^+^ Treg were captured by ψ-Angle (**Fig. 3h, 3i**). Thus, the ψ-Angle approach captured temporal changes of gene regulation as Timer protein matures.

Next, for the purpose of quantitative analysis, ψ-Angle data were categorised into the 5 Tocky loci (ψ-Tocky Locus, **Extended Data 4a**). ψ-Tocky Locus analysis allowed identification of differentially expressed genes for T cells in each ψ-Tocky Locus, revealing that immune checkpoint genes and activation/exhaustion-related genes were dynamically induced in T cells received TCR signals, and their expression was significantly changed across ψ-Tocky Loci (**Extended Data 4b – 4d**). In CD8 T cells, for example, *Eomes*, *Hspd1*, *Tnfrsf9*, *Bac*, and *Hprt* were induced in New and repressed in Persistent onwards; *Nfat1*, *Cd5*, *Slamf7*, *Zap70*, *Cd69* showed a peak in NPt; *Lck*, *CD8b1*, *Id2*, *CD3d*, *Il7r*, *Sell* (CD62L), *Rhoh*, and *Il7ra* were induced in the PAt and Arrested loci (**Extended Data 4b**). In Tconv, *Tnfrsf4* (OX-40) and the NFκB genes *Rel* and *Nfkb1* were induced in NPt, whilst the checkpoint *Icos* and *Ctla4*, Treg-associated genes *Tnfrsf18* (GITR) and *Ikzf2* (Helios), TGF-β receptor (*Tgfb1*) and IL2 receptor β chain (*Il2rb*) showed a peak in Persistent or PAt. Tconv cells in the Arrested locus showed increased *Sell*, *Bcl2*, and *Ccr7*, supporting that they were more enriched with less activated T cells (**Extended Data 4c**). In Treg, the TNFRSF molecules, including *Tnfrsf4, Tnfrsf9* (4-1BB), *and Tnfrsf18* and the NFκB genes *Rel*, *Nfkb1*, and *Nfkb2* were induced in either NPt or Persistent. The expression of the checkpoints *Pdcd1, Ctla4*, and *Icos* peaked at Persistent or PAt. Notably, Foxp3 expression was repressed in Persistent and increased in Arrested (**Extended Data 4d**). These data are compatible with the working model for eTreg that Foxp3 is controlled by TCR signalling, and that antigen-experienced Treg differentiate into eTreg, increasing the expression of Foxp3 and Treg-associated markers including CTLA-4 and ICOS ^41^.

In addition, ψ-Tocky Locus allowed pathway analysis for each stage of T cell activation following TCR signalling. Interestingly, RNA-related pathways were identified significantly enriched in genes upregulated during the New and NPt loci, while T cell activation-related pathways were enriched in genes upregulated at the Arrested locus (**Extended Data 4e** and **Fig. 3j, 3k**). This demonstrates that Nr4a3-Tocky successfully captures cellular activation processes that are promptly induced during the ultra-early phase of T cell activation, which has not been previously identified.

Lastly, we analysed the effects of aPD-L1 antibody on temporal changes in individual antigen-reactive T cells given their antigen engagement over time (**Fig. 3l**). Here we focused on perturbing the PD-1/PDa-L1 axis, as this is the commonest target across current clinical immunotherapy. Genes involved in the TCR signalling pathway and downstream, including *CD3e*, *Lat*, *Jun*, and *CD69* were mostly enhanced across ψ-Angle in Tconv cells, while *CD3d* and *Jun* were suppressed in CD8+ T cells (**Fig. 3l**). The expression of *Tox* and *Bhlhe40* showed a peak around 50° and 70°, respectively, whether in Tconv or Treg. The expression of *Il7r*, which is important for memory formation ^43^, was enhanced by aPD-L1 in CD8 and Tconv. Interestingly, aPD-L1 increased *Bcl2l11* expression in CD8 T cells as ψ-Angle increased, whilst enhancing *Bcl2* expression across ψ-Angle. Notably, *Ctla4* and *Icos* expression was enhanced by aPD-L1 in antigen-experienced Treg, with a peak around 75°, when the expression of *Malt1*, which is required for eTreg differentiation ^44^, also showed a peak.

In summary, the ψ-Angle approach using scRNA-seq data from Nr4a3-Tocky revealed the dynamics of the regulation of activation-induced genes and how these are modulated by aPD-L1, highlighting the reactive expression of the immune checkpoint molecules including *Ctla4*.

### Blockade of PD-L1 and CTLA-4 alters the temporal dynamics of TCR signalling of tumour-infiltrating T cells

The results above led to the hypothesis that the double blockade of PD-1 and CTLA-4 (standard of care for advanced melanoma), further changes the dynamics and immediate fate of tumour-reactive T cells. Here *Nr4a3*-tocky mice bearing B16-F10 melanoma tumours were treated with anti-CTLA-4 blocking antibody (aCTLA-4), anti-PD-L1 blocking antibody (aPD-L1), the combination of aCTLA-4 and aPD-L1 (aPD-L1/aCTLA-4), or their Isotype controls (**Extended Data 5a**). T cells in the tumour were analysed at Day 18 by flow cytometry.

The frequency of Timer^+^ antigen-reactive Treg was reduced by the administration of aPD-L1 and aPD-L1/aCTLA-4 (**Extended Data 5b, 5c**). Interestingly, the frequency of antigen-reactive Tconv was increased by aPD-L1/aCTLA-4 compared to the single therapies and the Isotype control (**Extended Data 5b, 5d**). The frequency of antigen-reactive CD8^+^ T cells showed a tendency to increase with aPD-L1/aCTLA-4 compared to the Isotype control and aCTLA-4 alone, although this did not reach statistical significance (p = 0.058) (**Extended Data 5b, 5e**). These results overall indicate that aPD-L1 and aCTLA-4 decreased the percentage of antigen-reactive Treg and increased the proportion of antigen-reactive Tconv and CD8^+^ T cells within the tumour. Thus, whilst Treg more dominantly interact with antigens in untreated tumours (Fig. 1), the combinatorial therapy made antigens within tumour more accessible to Tconv and CD8^+^ T-cells.

Interestingly, aPD-L1/aCTLA-4 reduced the Timer-Angle of Treg and CD8+ T-cells, but not Tconv, when compared to isotype control in tumour (**Extended Data 5f – 5h**), indicating that Treg and CD8+ T cells were more frequently engaged with antigen than Tconv on average and more susceptible to the effects of ICB. In addition, aPD-L1 alone also reduced the Timer-Angle of Treg, representing a decrease in Treg that are frequently engaging with antigen or progressing to the Arrested locus. This suggests that PD-L1 is driving the significant Treg effects seen with dual-checkpoint combination therapy.

The reduction of Timer-Angle in tumour-infiltrating Treg and CD8^+^ T-cells by aPD-L1/aCTLA-4 was largely due to the increase of New cells and the decrease of Arrested cells, in comparison to aCTLA-4 antibody alone and Isotype control (**Extended Data 5i, 5j**). The combinatorial therapy did not show any significant effects on the Tocky loci of Tconv (**Extended Data 5j**). In addition, aPD-L1/aCTLA-4 also increased the frequency of Treg cells in the NPt locus, suggesting that the increased Treg cells in the New locus survived and matured to increase those in the NPt locus but not further in terms of timer maturation (**Extended Data 5j**).

Collectively, dual aPDL-1/aCTLA-4 checkpoint blockade reduced antigen-reactive Treg and increased antigen-reactive Tconv and CD8^+^ T cells. Moreover, combination therapy accelerated the antigen recognition of Treg and CD8^+^ T cells, increasing T-cells in the New locus. However, this did not lead to the accumulation of T-cells with more mature Timer proteins, suggesting that they may have moved to other organs or died before Timer proteins matured.

### ICB reduces tumour-reactive eTreg and activates CD25^high^ Tconv and CD8 T cell populations

Next, we applied UMAP analysis to tumour-infiltrating T cell data in order to identify the tumour-reactive T cell populations that were controlled by blockade of PD-L1 and CTLA-4 at the single cell level (**Fig. 4**). Here we aimed to show which single-cell T-cell populations were increased or decreased by treatment, and test if these immunotherapies target any specific phase of TCR-induced activation.

**Figure 4.**
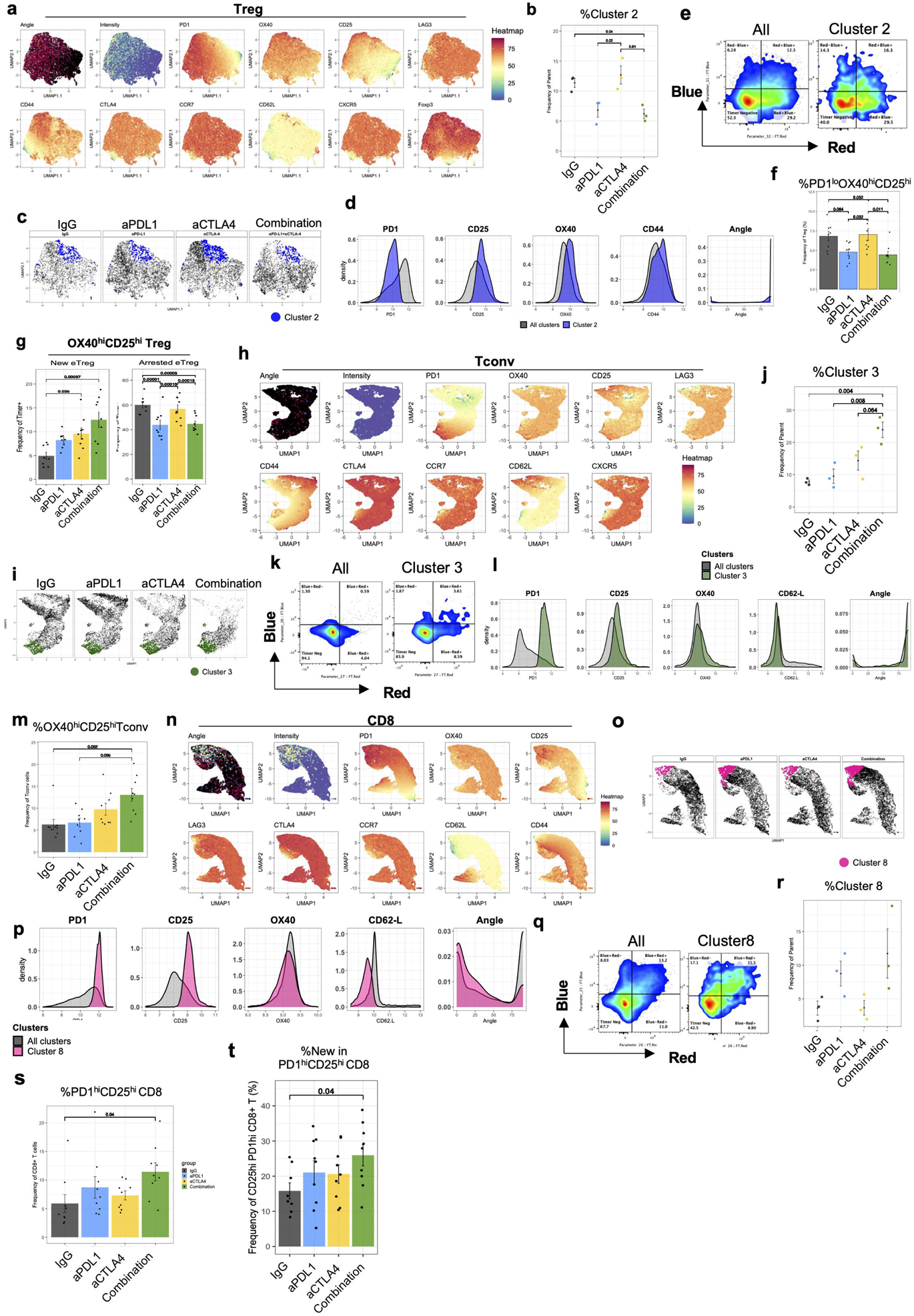
PD-1 and CTLA-4 blockade reduces tumour-reactive eTreg and activates CD25^high^ Tconv and CD8 T cell populations. Nr4a3-Tocky mice were treated with PD-L1 blocking antibody (aPD-L1), aCTLA-4 blocking antibody (aCTLA-4), the combination of aPD-L1 and aCTLA-4 (Combination), or Isotype Controls (IgG). Tumour-infiltrating T cells were analysed. **(a)** Heatmaps of UMAP plots showing Timer Angle, Intensity, and indicated markers in Treg from all the treatment groups. UMAP Analysis was performed using PD1, OX40, CD25, LAG3, CD44, extracellular CTLA4, CCR7, CD62L, CXCR5 and Foxp3. **(b)** Scatter plot showing the frequency of Treg in the UMAP cluster 2 by treatment groups. **(c)** Cluster 2 is highlighted by blue in the UMAP plot. **(d)** Overlaid histograms of indicated marker expression in Cluster 2 and all the clusters. **(e)** Representative flow cytometry plots showing Timer-Blue vs Timer-Red in Cluster 2 and all concatenated clusters (All clusters). (**f – g**) Bar plot showing the frequency of (**f**) PD1^lo^ CD25^hi^ OX40^hi^ Treg and (**g**) the percentage of CD25^hi^ OX40^hi^ Treg in the New (left) and Arrested locus (right)**. (h)** Heatmaps of UMAP plots showing Timer Angle, Intensity, and indicated markers in Tconv from all the treatment groups. UMAP Analysis was performed using PD1, OX40, CD25, LAG3, CD44, extracellular CTLA4, CCR7, CD62L and CXCR5. **(i)** Cluster 3 is highlighted by green in the UMAP plot. **(j)** Scatter plot showing the frequency of Cluster 3 by treatment groups. (**k**) Representative flow cytometry plots showing Blue vs Red in Cluster 3 and All clusters. **(l)** Overlaid histograms of indicated marker expression in Cluster 3 and All clusters. **(m)** Bar plot showing the frequency of CD25^hi^ OX40^hi^ Tconv **(n)** Heatmaps of UMAP plots showing Timer Angle, Intensity, and indicated markers in CD8+ T cells from all the treatment groups. UMAP Analysis was performed using PD1, OX40, CD25, LAG3, CD44, extracellular CTLA4, CCR7 and CD62L. (**o**) Cluster 8 is highlighted by red in the UMAP plot. **(p)** Overlaid histograms of indicated marker expression in Cluster 8 and All clusters. **(q)** Representative flow cytometry plots showing Blue vs Red in Cluster 8 and all clusters. **(r)**Scatter plot showing the frequency of Cluster 8 by treatment groups. **(s)** Bar plot showing the frequency of CD25^hi^ PD1 ^hi^ CD8+ T cells. **(t)** Bar plot showing the frequency of CD25^hi^ PD1^hi^ CD8+ T cells at the New locus. B16-melanoma bearing mice treated with either IgG, aPD-L1, aCTLA-4 and the combination of aPD-L1 and aCTLA-4 (n = 3 each). Data was collected over 3 independent experiments with a total of n=9 per group. Two-way ANOVA with Tukey’s multiple comparisons test was applied. Dots represent individual data points, error bars represent +/- standard error.

UMAP analysis of Treg showed that the expression of PD-1, CD25 and OX40 was associated with Angle and Intensity, indicating that these markers are expressed in relation to antigen-reactivity (**Fig. 4a**). Subsequently, computational clustering was applied to the UMAP result, identifying 8 clusters (**Extended Data 6a**). Cluster 2 was significantly reduced by both aPD-L1 alone and aPD-L1/aCTLA-4 combination therapy (**Fig. 4b, 4c**). The expression of CD25, OX40, and CD44 in Cluster 2 was significantly higher than the average, while PD-1 expression was significantly lower (**Fig. 4d**). Thus, Cluster 2 includes PD-1^lo^ CD25^hi^ OX40^hi^ CD44^high^ Tregs which were more antigen-reactive than the average (60% Timer^+^ in cluster 2 vs 47.7% in total Treg) (**Fig. 4e**).

The high expression of CD25 and OX40 is characteristic of eTreg ^40, 45^. We therefore established a new gating approach to identify the PD-1^lo^ CD25^hi^ OX40^hi^ eTreg across different datasets and tested if treatment depletes them (**Extended Data 6b**). In fact, the combination therapy as well as aPD-L1 alone reduced the percentages of PD-1^lo^CD25^hi^OX40^hi^ Treg and their parent population CD25^hi^OX40^hi^ Treg (**Fig. 4e, Extended Data 6c**). The combinatorial therapy reduced Timer^+^ cells in CD25^hi^OX40^hi^ eTreg and their mean Angle (**Extended Data 6d**). This was clearly due to the decline of CD25^hi^OX40^hi^ eTreg cells at the Arrested locus (**Extended Data 6e-6f**). On the other hand, CD25^hi^OX40^hi^ eTreg cells at the New locus were relatively preserved by the combination therapy, significantly increasing their proportion in the eTreg population (**Fig. 4g, Extended Data 6f**). These results suggest that the combination of aPD-L1 and aCTLA-4 did not prevent the antigen recognition of CD25^hi^OX40^hi^ eTreg but rather shortened the lifespan of this antigen-reactive Treg population.

Next, we analysed the effects of the ICB on tumour-infiltrating Tconv cells. UMAP analysis revealed an increase of Cluster 3 by aPD-L1/aCTLA-4 (**Fig. 4h – 4j, Extended Data 6g**). Tconv cells in Cluster 3 were more antigen reactive (14% in cluster3 and 5.9% in All), with increased cells in Blue^+^Red^+^ and Blue^-^Red^+^ quadrants (**Fig. 4k**). Tconv cells in Cluster3 were characteristically PD-1^hi^ OX40^hi^ CD25^hi^ CD62L^lo^ (**Fig. 4l**). Subsequently, we established a gating strategy to identify the CD25^hi^OX40^hi^ Tconv (**Extended Data 6h**), which were increased by the combination treatment (**Fig. 4m**), especially CD25^hi^OX40^hi^ Tconv within the NPt locus (**Extended Data 6i-6j**).

UMAP Analysis of tumour-infiltrating CD8^+^ T cells showed that combination treatment increased the frequency of cells in Cluster 8 (**Fig. 4n – 4r, Extended Data 6k**). CD8^+^ T cells in Cluster 8 were CD25^hi^ PD-1^hi^ OX40^int^ CD62L^lo^ (**Fig. 4p**) and were enriched with antigen-reactive CD8+ T cells (57.5% in cluster 8 vs 32.3% in All), which were mainly Timer-Blue^+^ (**Fig. 4q**) with an obviously higher proportion of New cells (**Fig. 4q**), indicating that these CD8^+^ T cells had recently encountered their cognate antigen. We established the gating of the CD25^hi^PD-1^hi^ CD8^+^ T cell population and confirmed the increase of this population by aPD-L1/aCTLA-4 (**Extended Data 6l, Fig. 4s**). The combination treatment increased CD25^hi^PD-1^hi^ CD8^+^ T cells at the New locus (**Fig. 4t**), although these T cells did not accumulated in more timer mature loci (**Extended Data 6j-6k**).

These results overall suggest that the combination of PD-L1 and CTLA-4 blocking antibodies specifically depleted PD-1^lo^ CD25^hi^ OX40^hi^ CD44^high^ and CD25^hi^OX40^hi^ eTreg populations, whilst CD25^hi^ PD-1^hi^ OX40^int^ CD62L^lo^ and CD25^hi^PD-1^hi^ CD8^+^ T cell populations and PD-1^hi^ OX40^hi^ CD25^hi^ CD62L^lo^ and CD25^hi^OX40^hi^ Tconv populations instead gained more access to their cognate antigen, and increased in number.

### Addition of an OX40 agonist enhances the effects of the combination checkpoint blockade on tumour-reactive T cells in tumour-draining lymph nodes

Having elucidated the effects of PD-L1 and CTLA-4 blocking antibodies on the OX40^hi^ T cell populations, we hypothesized that the OX40 agonist OX-86 will enhance the effects of the combination therapy on tumour-reactive T cells (**Fig. 5**). Firstly, we investigated the effects of the triple therapy on tumour-reactive T cells in tdLN, given the intriguing effects of the double combination of tumour-reactive CD4 and CD8 T cells in tdLN as demonstrated above.

**Figure 5:**
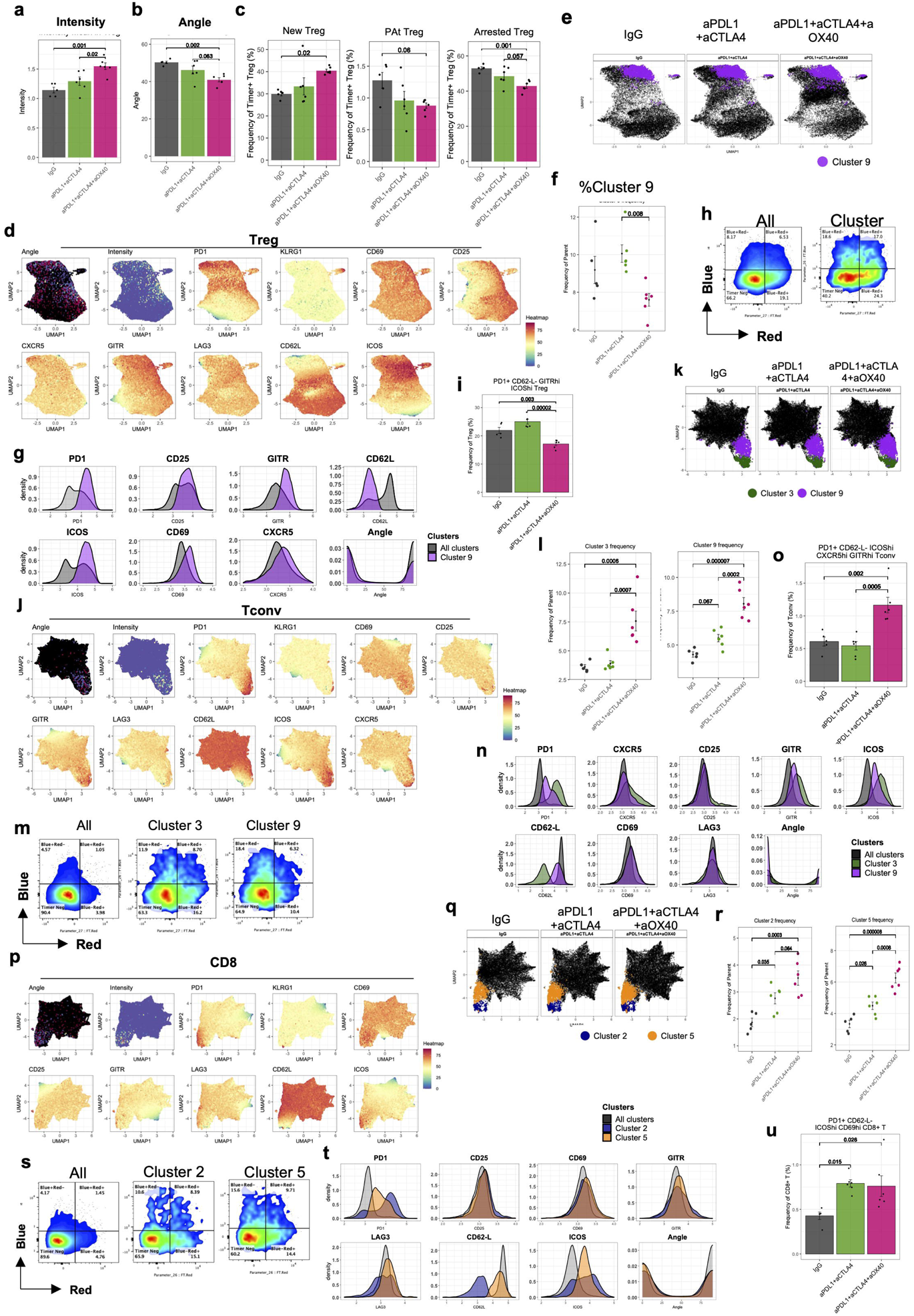
Addition of OX40 agonist to the combination enhances tumour-reactive effector T cell activities and decreases tumour-reactive Treg. B16-melanoma bearing Nr4a3-Tocky mice were treated with the double combination of aPD-L1 and aCTLA-4, the triple combination of aPD-L1, aCTLA-4 and aOX40 and Isotype Controls (IgG) (n=6 each, aCTLA-4 treatments on days 12, 14, and 16; aPD-L1 and aOX40 treatments on days 14 and 16) and T cells from tumour-draining lymph nodes (tdLN) were analysed on d18. (**a-c**) Bar plots showing (**a**) Intensity, (**b**) Angle, (**c**) the percentage of cells in each of the New, Pat, and Arrested loci. **(d)** Heatmaps of UMAP plots showing Timer Angle, Intensity, and indicated markers in Treg from all the treatment groups. UMAP Analysis was performed using PD1, KLRG1, CD69, CD25, CXCR5, GITR, LAG3, CD62L and ICOS. **(e)**Cluster 9 is highlighted by purple in the UMAP plot. **(f)** Scatter plot showing the frequency of Cluster 9 by treatment groups. **(g)** Overlaid histograms of indicated marker expression in Custer 9 and all clusters concatenated (All clusters). **(h)** Representative flow cytometry plots showing Blue vs Red in Cluster 9 and All clusters. **(i)** Bar plot showing the frequency of PD1^hi^ CD62L^-^ GITR^hi^ ICOS^hi^ Treg out of the tdLN Treg population. **(j)** Heatmaps of UMAP plots showing Timer Angle, Intensity, and indicated markers in Treg from all the treatment groups. UMAP Analysis was performed using PD1, KLRG1, CD69, CD25, CXCR5, GITR, LAG3, CD62L and ICOS. **(k)** Clusters 3 and 9 are highlighted in the UMAP plot. **(l)** Scatter plot showing the frequency of Clusters 3 (left) and 9 (right) by treatment groups. **(m)** Representative flow cytometry plots showing Blue vs Red in Clusters 3 and 9 and All clusters. **(n)** Overlaid histograms of indicated marker expression in Clusters 3 and 9 and All clusters. **(o)** Bar plot showing the frequency of PD1^hi^ CD62L^-^ ICOS^hi^ CXCR5^hi^ GITR^hi^ Tconv. **(p)** Heatmaps of UMAP plots showing Timer Angle, Intensity, and indicated markers in Treg from all the treatment groups. UMAP Analysis was performed using PD1, KLRG1, CD69, CD25, GITR, LAG3, CD62L and ICOS. **(q)** Clusters 2 and 5 are highlighted in the UMAP plot. **(r)**Scatter plots showing the frequency of CD8+ T cells in Clusters 2 (left) and 5 (right) by treatment groups. **(s)** Representative flow cytometry plots showing *Blue* vs *Nr4a3*-Red in clusters 2 and 5 and all concatenated clusters. **(t)** Overlaid histograms of indicated marker expression in Clusters 2 and 5 and All clusters. (**u**) Bar plot showing the frequency of PD1^hi^ CD62L^-^ ICOS^hi^ CD69^hi^ CD8+ T cells by treatment groups. Data were collected over an experiment with a total of n=6 per group. One-way ANOVA with Tukey’s multiple comparisons test was applied. Dots represent individual data points, error bars represent +/- standard error.

The triple combination therapy increased Timer Intensity and decreased Angle in Treg from tdLN, in comparison to the isotype control and the double combination, indicating a longer and more frequent interaction with antigen (**Fig. 5a, 5b**). The triple combination increased Treg cells in the New locus, decreasing those in the PAt and Arrested loci (**Fig. 5c and Extended Data 7a-7b**), suggesting a shorter life for them after cognate antigen signalling. UMAP analysis of tdLN Tregs showed that the triple combination decreased the percentage of Cluster 9 in comparison to the double combination (**Extended Data 7c, Fig. 5d – 5f**). Treg in Cluster 9 highly expressed PD-1, GITR, ICOS and CD69 while showing a low CD62L expression (**Fig. 5g**). Cells in Cluster 9 were enriched with antigen-reactive Treg (60% timer positive in Cluster 9 and 35% in All, **Fig. 5h**) and had a high Timer Intensity (**Extended Data 7d**), indicating that Cluster-9 Treg had received significant TCR signals over time. Interestingly, this cell population shares common features with the anti-tumour Treg population shown in Figure 2. In fact, the triple combination significantly decreased PD-1^hi^ CD62L^lo^ ICOS^hi^ GITR^hi^ Treg and their parent population PD-1^hi^ CD62L^lo^ Treg (**Fig. 5i**, **Extended Data 7e-7f**).

UMAP analysis of Tconv in tdLN showed the triple combination increased cells in Clusters 3 and 9 (**Fig. 5j – 5l**), which were enriched with Timer+ antigen-reactive T cells, with more mature cells in Cluster 9 than Cluster 3 (**Fig. 5m**, **Extended Data 7h**), suggesting that Cluster 9 cells differentiate into Cluster 3 cells. Cluster-3 Tconv were PD-1^hi^ CXCR5^hi^ CD25^hi^ GITR^hi^ ICOS^hi^ CD69^hi^ CD62L ^lo^ (**Fig. 5n**), a profile similar to Tfh cells as well as to the tumour-reactive PD-1^hi^ ICOS^hi^ CXCR5^hi^ GITR^hi^ CD62L^-^ Tconv in Figure 2. Interestingly, this Tfh-like population was further increased by the triple combination immunotherapy aPD-L1/aCTLA-4/aOX40 (**Fig. 5o, Extended Data 7i**). A Parent population of the Tconv population, PD-1^hi^ CD62L^-^ Tconv was also uniquely increased in the triple combination treatment group (**Extended Data 7j**). Cells in Cluster 9 had intermediate expression of PD-1, CXCR5, GITR, ICOS and LAG3, and high expression of CD62L and CD69 (**Fig. 5n**). These results suggest that the triple combination immunotherapy further increased the frequency of anti-tumour PD-1^hi^ CD62L^-^ Tconv populations, including fully mature PD-1^hi^ ICOS^hi^ CXCR5^hi^ GITR^hi^ CD62L^-^ Tconv cells, and the transitional population PD-1^int^ ICOS ^int^ CXCR5^int^ GITR^int^ CD62L^-^ Tconv cells.

Next, UMAP analysis of tdLN CD8^+^ T cells showed that the triple combination increased the percentages of CD8^+^ T cells in Clusters 2 and 5 (**Fig. 5p-7r, Extended Data 7k**). Both clusters were enriched with Timer^+^ antigen-reactive T cells (**Fig. 5s**), showing the lowest Angle and highest intensity, indicating a stronger and more frequent antigen recognition (**Extended Data 7l**). CD8^+^ T cells in Cluster 2 were PD-1^hi^ CD62L^-^ ICOS^hi^ CD69^hi^, showing the features of the tumour reactive CD8^+^ T cell population in the steady-state tdDN (Figure 2, **Fig. 5t**). This CD8^+^ T cell population and its parent population PD-1^hi^ CD62L^-^ CD8^+^ T cells, were further increased in the tdLN by the triple combination (**Fig. 5u, Extended Data 7m**). CD8^+^ T cells in Cluster 5 were PD-1^int^ CD62L^-^ ICOS^int^ CD69^hi^, suggesting that they are a transitioning cell population, maturing towards the cells in Cluster 2 (**Extended Data 7n**).

Collectively, these results suggest that the triple combination aPD-L1/aCTLA-4/aOX40 immunotherapy further enhanced the activation of tumour-reactive T cells in tdLN. Tumour-reactive PD-1^hi^ CD62L^lo^ Treg were reduced, most probably through accelerating the death of anti-tumour Treg population. Meanwhile, tumour-reactive PD-1^hi^ CD62L^-^ T conv and PD-1^hi^ CD62L^-^ ICOS^hi^ CD69^hi^ CD8+ T cells were increased upon the triple combination.

### The OX40 agonist enhances the effects of the combination checkpoint blockade on the tumour-infiltrating, reactive T cells in a unique manner

Lastly, we investigated the effects of the triple combination therapy on tumour-infiltrating T cells. Similar to the double combination aCTLA-4 and aPD-L1, the triple combination with aOX40 decreased Angle (**Fig. 6a-6c**), increased cells at the New locus and reduced those at the Arrested locus (**Fig. 6d**). UMAP analysis identified 9 clusters (**Fig. 6e, Extended Data 8a)**. Cluster 7 had the highest Angle and Intensity (**Fig. 6f, Extended Data 8b**), and the triple combination showed a trend towards decreasing this population relative to double therapy, although this was not statistically significant (**Fig. 6g**). Treg in Cluster 7 were PD-1^hi^ GITR^hi^ ICOS^hi^ LAG3^hi^ CD25 ^intermediate^ CD69^low^, with 80% Timer+, mostly in the Blue+Red+ and Blue- Red+ quadrants (**Fig. 6h – 6i**).

**Figure 6:**
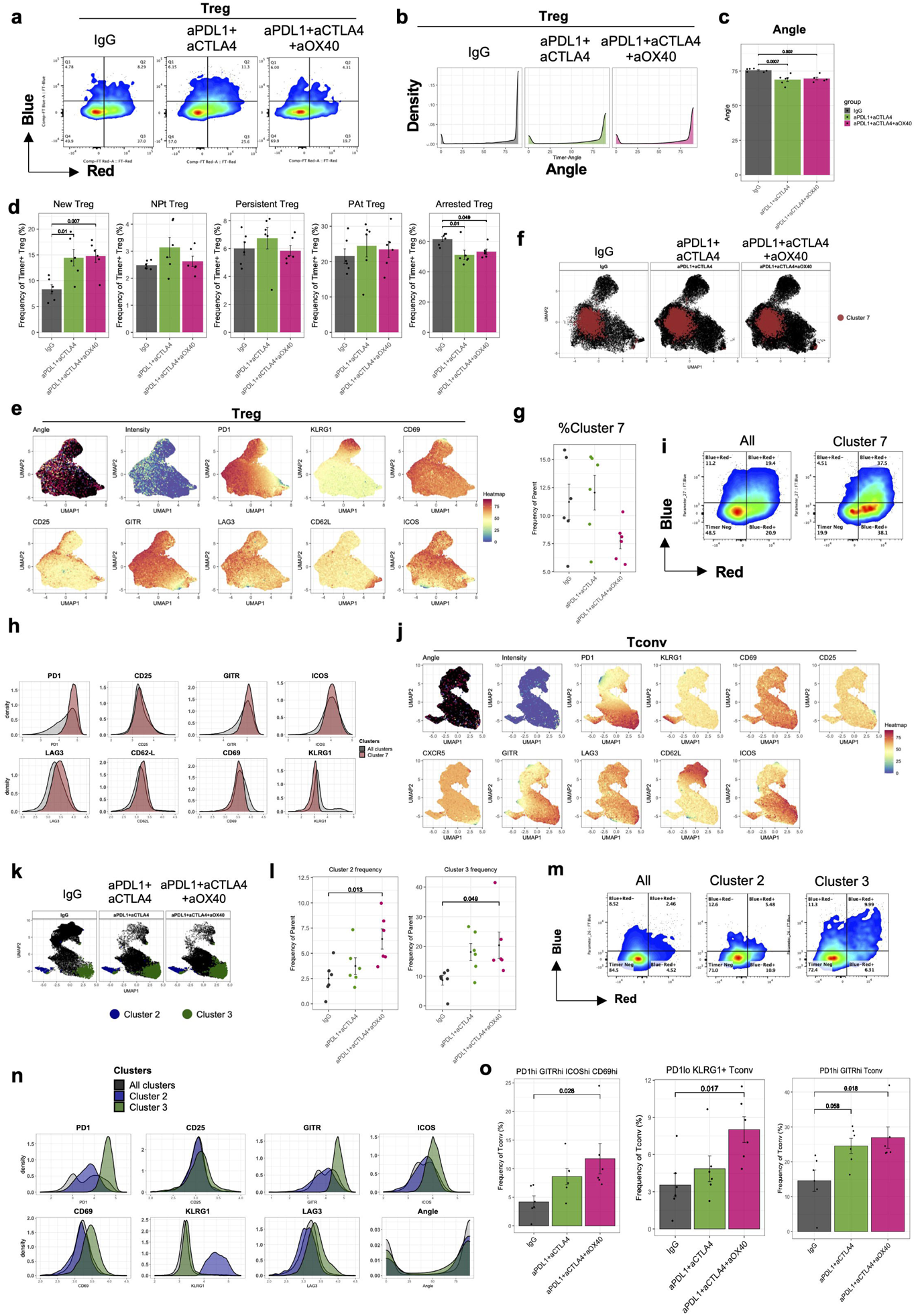
Addition of an OX40 agonist to the combination changes the early fate of tumour infiltrating CD4+ T cells. Nr4a3-Tocky mice treated from B16-melanoma bearing mice were treated with the double combination of aPD-L1 and aCTLA-4, the triple combination of aPD- L1, aCTLA-4 and aOX40 and Isotype Controls (IgG) (n=6 per group). **(a)** Representative flow cytometry plots of Tumour-infiltrating Treg. **(b)** Density plot showing Angle distribution in Treg from each group. **(c)** Bar plot showing mean Angle. **(d)** Bar plots showing the frequency of Treg cells in each Tocky Locus out of Timer+ Treg. **(e)**Heatmaps of Angle, Intensity, and indicated markers in a UMAP plot, which was calculated by PD1, KLRG1, CD69, CD25, GITR, LAG3, CD62L and ICOS. **(f)** Cluster 7 is highlighted in the UMAP plot. **(g)** Scatter plots showing the frequency of Treg in Cluster 7. **(h)** Overlaid histograms of indicated marker expression in Cluster 7 and all clusters concatenated (All). **(i)** Representative flow cytometry plots of Cluster 7 and All. **(j)** Heatmaps of Angle, Intensity, and indicated markers in a UMAP plot, which was calculated by PD1, KLRG1, CD69, CD25, GITR, LAG3, CD62L and ICOS. **(k)** Clusters 2 and 3 are highlighted in the UMAP plot. **(l)** Scatter plots showing the frequency of Tconv in Clusters 2 and 3. **(m)** Representative flow cytometry plots of Clusters 2 and 3 and All. **(n)** Overlaid histograms of indicated marker expression in Clusters 2 and 3 and All. **(o)** Bar plots showing the frequency of indicated populations. Dots represent individual data points, error bars represent +/- standard error. Thresholds for Blue and Red were set using a fully stained WT mouse.

UMAP analysis of tumour-infiltrating Tconv identified 9 clusters (**Fig. 6j**, **Extended Data 8c**) and showed that the triple combination significantly increased Tconv cells in Clusters 2 and 3 only (**Fig. 6k – 6l**). Cluster 3 Tconv were the most antigen reactive compared to all clusters (30% vs 15%), and were enriched with Blue+Red- or Blue-Red+ cells, but not Blue+Red+ cells (**Fig. 6m**); this Timer profile indicates that they had received short (not persistent) cognate antigen signals ^28^. Tconv in Cluster 2 had low to intermediate expression of PD-1, CD25, GITR, ICOS, CD69 and LAG3, while showing characteristically high KLRG1 expression. In contrast, Tconv in Cluster 3 were PD-1^hi^ CD25^hi^ GITR^hi^ ICOS^hi^ CD69^hi^ LAG3^hi^ with a low KLRG1 expression (**Fig. 6n**). Interestingly, Tconv in Cluster 3 had an abundant Blue+Red+ quadrant, indicating their frequent antigen recognition (**Fig. 6n**). In fact, the triple combination immunotherapy increased the logically-gated PD-1^low^ KLRG1^+^ Tconv (Cluster 2 equivalent), PD-1^hi^ GITR^hi^ ICOS^hi^ CD69^hi^ Tconv (Cluster 3 equivalent) and their parent population PD-1^hi^ GITR^hi^ Tconv (**Extended Data 8e-8f, Fig. 6o**).

Finally, tumour-infiltrating CD8^+^ T cells showed that the triple combination increased CD8+ T cells in the New locus and decreased those in the Arrested locus, decreasing mean Angle (**Fig. 7a – 7d**). UMAP analysis identified 9 clusters (**Fig. 7e, Extended Data 9a**) and Clusters 4 and 8 only were significantly increased by the triple combination (**Fig. 7f -7g**). Indeed, CD8+ T cells in Cluster 4 were PD-1^hi^ CD25^hi^ CD69^hi^ GITR^hi^ LAG3^hi^, while showing CD62L^low^ KLRG ^low^ (**Fig. 7h**). CD8+ T cells in Cluster 8 had intermediate to low expression levels of CD25, CD69, GITR, ICOS, and LAG3, while showing high KLRG1 expression. Nearly half of CD8+ T cells in Cluster 4 were found in the Blue+Red+ quadrant in a diagonal pattern, highlighting that this population persistently recognised antigen, whilst CD8+ T cells in Cluster 4 had relatively more New cells (Blue+Red-), indicating that they did not survive long or moved to other tissues after receiving TCR signals (**Fig. 7i**). A logical gating approach (**Extended Data 9b-9c)** showed that the triple combination therapy increased the Cluster 4 equivalent PD-1^hi^ GITR^hi^ CD69^hi^ CD25^hi^ KLRG1^-^ CD62L^-^, their parent PD-1^hi^ GITR^hi^ CD69^hi^ CD25^hi^ (**Fig. 7j**) the Cluster 8 equivalent PD-1^hi^ CD62L^-^ KLRG1^+^, and their parent PD-1^hi^ CD62L- CD8^+^ T cell populations - (**Fig. 7k**).

**Figure 7:**
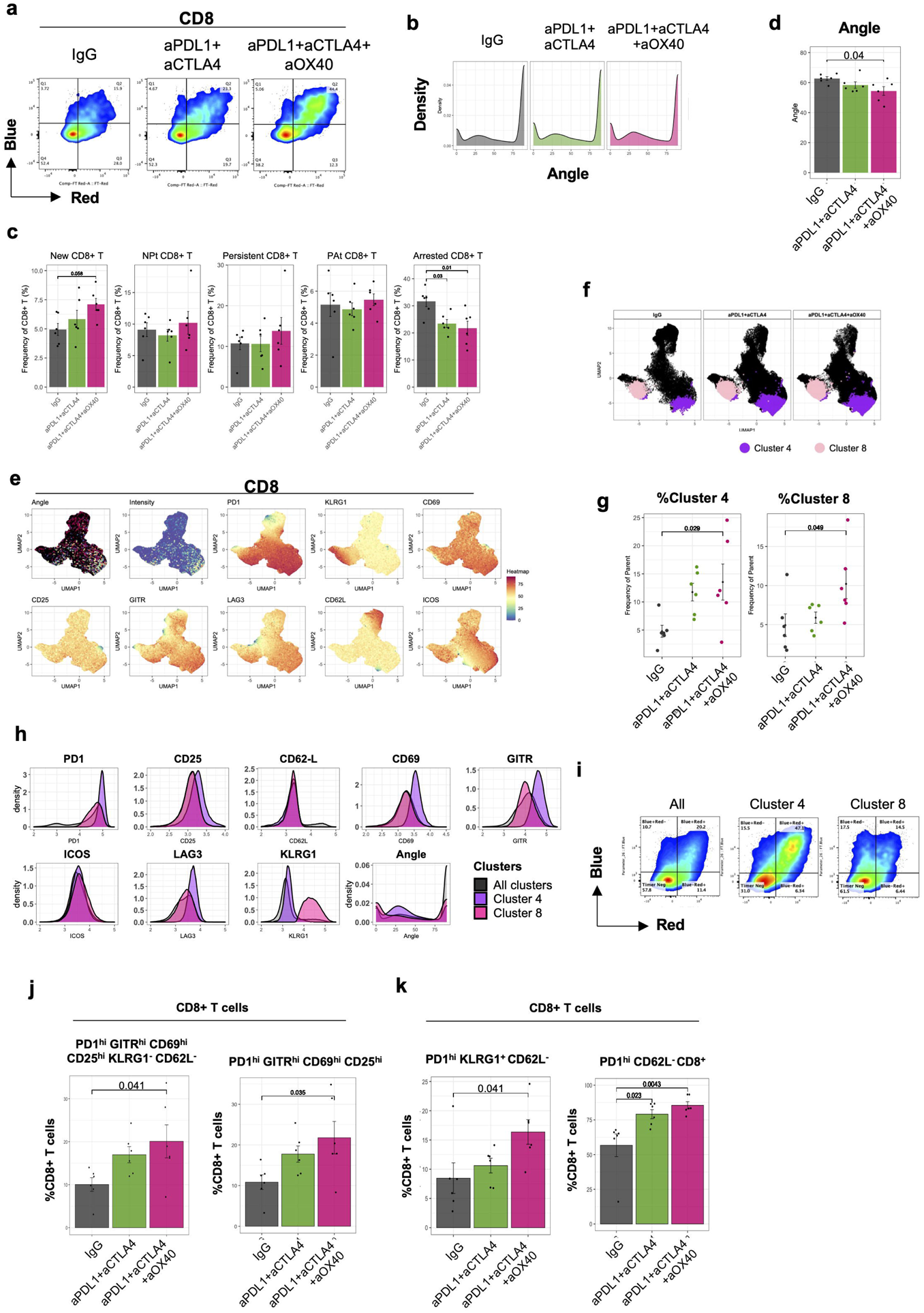
Addition of an OX40 agonist to the combination changes the early fate of tumour infiltrating CD8+ T cells. The same analysis in Fig. 6 was performed for CD8+ T cell. (**a**) Representative flow cytometry plots of tumour-infiltrating CD8+ T cells. **(b)** Density plot showing Angle distribution in Treg from each group. **(c)** Bar plots showing the frequency of Treg cells in each Tocky Locus out of Timer+ Treg. **(d)** Bar plot showing mean Angle. (**e**) Heatmaps of Angle, Intensity, and indicated markers in a UMAP plot, which was calculated by PD1, KLRG1, CD69, CD25, GITR, LAG3, CD62L and ICOS. **(f)** Clusters 4 and 8 are highlighted in the UMAP plot. **(g)** Scatter plots showing the frequency of CD8+ T cells in Clusters 4 and 8. **(h)** Overlaid histograms of indicated marker expression in Clusters 4 and 8 and all clusters concatenated (All). **(i)** Representative flow cytometry plots of Clusters 4, 8 and All. (**j-k**) Bar plots showing the frequency of indicated populations. One-way ANOVA with Tukey’s multiple comparisons test was applied. Dots represent individual data points, error bars represent +/- standard error.

Collectively, the addition of aOX40 to the double combination therapy aPD-L1 and aCTLA-4 enhanced the effects of the double combination therapy, reactivating and expanding not only pre-existing tumour-reactive T cells but also transitional effector T cell populations in both TILs and tdLN. Importantly, the triple combination decreased tumour-reactive PD-1^hi^ Treg populations, increasing highly antigen-reactive effector-like CD8 and Tconv populations and KLRG1 expressing Tconv and CD8+ T cell populations.

## Discussion

By analysing temporal profiles of individual antigen-reactive T cells in vivo, the current study revealed that PD-1 and CTLA-4 blockade depleted OX40 ^hi^CD25^hi^ eTreg while promoting the differentiation of OX40 ^hi^CD25^hi^ Tconv and two distinct CD8 Tocky populations, KLRG1^hi^PD-1^int^ and KLRG1^lo^PD-1^hi^ CD8 T cells. These effects were further enhanced by the addition of aOX40 to the combinatorial therapy. KLRG1^hi^PD-1^int^ CD8 T cells were remarkably more expanded in tumour than tdLN, showing relatively higher percentage of cells in the New locus. This suggests that KLRG1^hi^PD-1^int^ CD8 T cells received short TCR signalling and most probably prematurely died in tumour. On the other hand, the majority of KLRG1^lo^PD-1^hi^ CD8 T cells in the tumour were Timer+, mainly in the Persistent locus.

The apparent short-life of KLRG1^hi^PD-1^int^ CD8 T cells by Tocky analysis is compatible with the evidence that KLRG1 is a marker for short-lived effector CD8 T cells in viral infections ^46^ and that effector CD8+ T cells acquire KLRG1 expression when infiltrating tissues ^47^. In cancer, the combination of aOX40 and anti-4-1BB increases KLRG1^hi^PD-1^int^ CD8 T cells in M38-bearing mice ^48^. On the other hand, the high frequency of KLRG1^lo^PD-1^hi^ CD8 T cells at the Persistent locus implies that these T cells are found in both tdLN and tumour, actively interacting with antigen recognition presenting cells. These suggest that KLRG1^lo^PD-1^hi^ CD8 T cells are in the phase of priming in tdLN (hence persistent TCR signalling), and that KLRG1^lo^PD-1^hi^ CD8 cells can generate terminally differentiated KLRG1^hi^PD-1^int^ CD8 T cells which migrate to the tumour, where they no longer interact with APCs and function as cytotoxic T cells (hence shorter TCR signalling only for killing). These TCR signal dynamics are compatible with the microscopic finding that the contact between a cytotoxic T cell and its target cell lasts for several minutes only ^49^, while a cognate CD8 T cell and a dendritic cell interact for several hours in CD8 T cell priming ^50^. Alternatively, Some KLRG1+ CD8+ T cells may lose KLRG1 expression in a Bach2 dependent manner and differentiate into memory T cells ^51^. It is of interest to investigate if clinical effects are correlated with either or both of KLRG1^lo^PD-1^hi^ CD8 T cells and KLRG1^hi^PD-1^int^ CD8 T cells. Furthermore, investigations using TCR-seq will reveal the lineage relationship between KLRG1^hi^PD-1^int^ and KLRG1^lo^PD-1^hi^ in future studies.

Interestingly, the combined blockade of PD-L1 and CTLA-4 specifically reduced eTreg. Some studies using checkpoint blockade showed the expansion or activation of Treg. For example, Marangoni et al showed that CTLA-4 blockade induced CD28-mediated expansion of Treg ^52^. In addition, antibody-dependent cellular cytotoxicity may deplete Treg upon CTLA-4 blockade ^53^. The effects of checkpoint blockade on the eTreg was further enhanced by the inclusion of aOX40 in the current study, which replicates our previous finding that the anti-OX40 antibody depleted eTreg, which were identified as persistent Foxp3 expressors using Foxp3-Tocky, in a skin allergy model ^40^. OX40 signalling is considered to be anti-apoptotic, mainly based on the report that in vitro activation of OX40^-/-^ T cells results in accelerated apoptosis due to low expression of anti-apoptotic Bcl2 ^54^. Possible mechanisms for this Treg depletion includes activation-induced cell death through Bim-mediated apoptosis, or a relative deprivation of cytokines, especially IL-2, leading to a failure to meet their metabolism demands in the tumour microenvironment. Future studies are needed to address these questions.

The ICB increased the antigen-reactive Tconv that highly expressed immune checkpoints OX40, PD1 or ICOS, and the Treg-associated surface molecules CD25 and GITR. These results lead to the hypothesis that tumour-reactive Tconv and eTreg compete for not only antigen, but also costimulatory TNFRSF ligands (OX40-L, ICOS-L, and GITR-L), and cytokines (CD25 for IL-2). PD-1 blockade through aPD-L1 antibody releases PD-1-mediated inhibitory signalling in T cells ^32^. The addition of aOX40 to the double combination further enhanced the antigen-reactive Tconv populations PD1^hi^ CD62L^-^ and PD1^hi^ CD62L^-^ ICOS^hi^ CXCR5^hi^ GITR^hi^, which suggests that aOX40 further boosted and expanded the expansion of PD1^hi^ population. This resonates with the findings that the agonistic aOX40 antibody expands PD1^hi^ CXCR5^hi^ Tfh cells in a *plasmodium* infection model ^55^ and the differentiation of PD1^hi^ CXCR5^hi^ BCL6^hi^ Tfh-like cells in human melanoma tissues ^38^. However, the effect of the triple combinatorial therapy is not specific to the Tfh population, since the PD1^hi^ CD62L^-^ ICOS^hi^ CXCR5^hi^ GITR^hi^ Tconv is phenotypically similar to TFR and activated Treg, apart from their Foxp3 expression. Notably, the Tconv population have high GITR expression, which is typically specific to Treg^56^. However, it is unlikely that they immediately differentiate into Treg given their low CD25 expression, which mediates the IL-2 signalling that is important for sustaining Foxp3 transcription ^40^. It is plausible that checkpoint blockade allows Tconv to compete with Treg for the binding to TNRSF and other ligands on APCs, a hypothesis best tested for each of antigen-specific Tconv and Treg using TCR sequencing analysis.

The multidimensional Tocky scRNA-seq analysis revealed hidden dynamics of gene expression in tumour-reactive T cells and how they are manipulated by the immunotherapy. The ψ-Angle analysis showed that the expression of *Ctla4* and *Icos* was induced in antigen-experienced Treg specifically in their late activation phase (i.e. high ψ-Angle) by aPD-L1. This means that aPD-L1 mediated its effects on Treg only when Treg are activated after engaging with antigen for some time. Interestingly, antigen-reactive CD8+ T cells upregulated the pro-apoptotic gene *Bcl2l11* (Bim) in the presence of aPD-L1, which also induced the anti-apoptotic gene *Bcl2* expression as well, while the two genes were not identified as DEG for aPD-L1 in CD4+ T cells. This suggests that aPD-L1 specifically targets apoptotic mechanism in CD8+ T cells, although further studies are needed to investigate tumour-infiltrating T cells. In Tconv, aPD-L1 induced activation-related genes *Bhlhe40*, *Lat*, and *Tox* specifically in Tconv in the Persistent locus. This suggests that aPD-L1 has unique and specific effects on tumour-reactive Tconv. Studies using T-cell-specific *Bhlhe40* KO mic showed that Bhlhe40 plays roles in cytokine regulation in Tconv ^57^. A recent report showed that T cells required *Bhlhe40* to mediate the effects of the combination of aPD-L1 and aCTLA-4 on the rejection of a sarcoma line ^58^. The role of Bhlhe40 in antigen-reactive Treg is not known. Tox play roles in CD8 T cell exhaustion ^59^ but its role in CD4 T cells is unclear. It is expected that the significance of the gene signature of each Tocky population will become clearer by an extensive study correlating clinical effects of each immunotherapy and single cell Tconv and CD8 T cell populations.

Whilst many clinical investigations and trials for combinatorial checkpoint therapies are on-going worldwide, it is hoped that basic studies can elucidate mechanisms underlying each checkpoint blockade ^22, 60^. To this end, the current study used single cell technologies, revealing the time dimension of tumour-reactive T cells and their dynamic response to checkpoint blockade therapies, providing insights into single-cell level mechanisms of checkpoint blockade in the cancer context.

## Acknowledgements

MO was supported by a CRUK Programme Foundation Award and the MRC grant (MR/S000208/1). AM was supported by a CRUK Programme Award (CRM183X) and the Institute of Cancer Research/Royal Marsden Hospital Centre for Translational Immunotherapy. KH was supported by CRUK and the Myfanwy Townsend Melanoma Research Fund. JH was supported by an ICR studentship from CRUK Convergence Science Center. This research was also supported by KAKENHI research grants from the Japan Society for the Promotion of Science (JSPS) (JP19H05426, JP21K07082, and JP21H00433, to MO; JP18H05417 and JP22H00448 to TO; JP22H02888 to IO) and Japan Agency for Medical Research and Development (grant numbers JP21jm0210074 to YS and MO). We would like to thank Dr Jessica Rowley and Dr Larissa Zárate-García (Flow Cytometry Facility, South Kensington, Imperial College London) and Dr Shintaro Funasaki (Kumamoto University) for supporting flow cytometric analysis and sorting, and the Central Biomedical Services (Imperial College London, Institute of Cancer Research) and the Center for Animal Resources and Development (Kumamoto University) for supporting animal experiments.

