## Extended Data Legends for "Single-cell level temporal profiling of tumour-reactive T cells under immune checkpoint blockade"

**Extended Data Fig. 1: *Nr4a3* tocky reveals Tumour-specific effects of temporal dynamics of TCR signalling in main T cell populations.**

a- Barplots representing the frequency of CD8+ T cells in different TockyLoci out of the parent CD8+ T cell population. Data was collected over 3 independent experiments with a total of n=9 per group.Two-way ANOVA with Tukey's multiple comparisons test was applied. Dots represent individual data points, error bars represent +/- standard error.

b- Barplots representing the frequency of Tconv cells in different TockyLoci out of the parent Tconv cell population. Data was collected over 3 independent experiments with a total of n=9 per group.Two-way ANOVA with Tukey's multiple comparisons test was applied. Dots represent individual data points, error bars represent +/- standard error.

c- Barplots representing the frequency of Treg cells in different TockyLoci out of the parent Treg cell population. Data was collected over 3 independent experiments with a total of n=9 per group.Two-way ANOVA with Tukey's multiple comparisons test was applied. Dots represent individual data points, error bars represent +/- standard error.

**Extended Data Fig. 2: Identification of tumour-reactive T-cell populations by *Nr4a3*-Tocky**

a- Representation of the clusters identified by k-means on the PCA result of parameters selected for UMAP in the UMAP space of cLN and tdLN Treg (n=5).

b- Representation of the clusters identified by k-means on the PCA result of parameters selected for UMAP in the UMAP space of cLN and tdLN Tconv (n=5).

c- Representation of the clusters identified by k-means on the PCA result of parameters selected for UMAP in the UMAP space of cLN and tdLN CD8+ T cells (n=5).

d- Representative flow cytometry plots of the gating strategy to identify Tconv populations induced by tumour burden in the cLN and tdLN, this includes PD1+ CD62L- ICOShi CXCR5hi GITRhi cells and their Nr4a3-Blue vs Red pattern.

e- Representative flow cytometry plots of the gating strategy to identify Treg populations induced by tumour burden in the cLN and tdLN, this includes PD1+ CD62L- GITRhi ICOShi OX40hi cells and their Nr4a3-Blue vs Red pattern.

f- Representative flow cytometry plots of the gating strategy to identify CD8+ T populations induced by tumour burden in the cLN and tdLN, this includes PD1+ CD62L- ICOShi CD69hi cells and their Nr4a3-Blue vs Red pattern.

g- Barplot representing the frequency of PD1+ CD62L- Tconv in either the cLN or tdLN of B16-F10 tumour bearing mice (n=5). Student's t-test, dots represent individual data points, error bars represent +/- standard error.

h- Barplot representing the frequency of PD1+ CD62L- Treg in either the cLN or tdLN of B16-F10 tumour bearing mice (n=5). Student's t-test, dots represent individual data points, error bars represent +/- standard error.

i- Barplot representing the frequency of PD1+ CD62L- CD8 T cells in either the cLN or tdLN of B16-F10 tumour bearing mice (n=5). Student’s t-test, dots represent individual data points, error bars represent +/- standard error.

j- Visual representation of Pearson Correlation analysis of PD1, ICOS and CD69 expression and Timer Intensity in CD8+ T cells in cLN and tdLN concatenated (n=5). Each dot represents an individual cell.

**Extended Data 3: TCR signalling dynamically controls the expression of immune checkpoint and costimulatory molecules on tumour-infiltrating T cells.**

(a) Scatter plots showing normalised mean fluorescence intensity (MFI) of PD-1, CD25, OX40, GITR, ICOS and LAG3 across Tocky loci in tumour infiltrating Treg.

(b) Visual representation of Pearson Correlation analysis of Timer Intensity vs PD-1, CD25, OX40, GITR, ICOS and LAG3 expression in tumour-infiltrating Treg.

(c) Scatter plots showing normalised MFI of PD-1, OX40, LAG3 and ICOS across Tocky loci in Tumour infiltrating Tconv.

(d) Visual representation of Pearson Correlation analysis of Timer Intensity vs PD1, OX40, LAG3 and ICOS expression in tumour-infiltrating Tconv.

(e) Scatter plots showing normalised MFI of OX40, PD1, CD62L, LAG3, CD25, ICOS and GITR across Tocky loci in tumour infiltrating CD8+ T cells.

(f) Visual representation of Pearson Correlation analysis of Timer Intensity vs OX40, PD1, CD62L, LAG3, CD25, ICOS and GITR expression in tumour-infiltrating CD8+ T cells.

Representative data of four independent experiments (n=6 represented). Each dot represents an individual cell. Two-way ANOVA with Tukey’s multiple comparisons test was applied. Dots represent individual data points, error bars represent +/- standard error. Data was collected over 4 independent experiments with a total of n=15 per group.

**Extended Data 4: Multidimensional Tocky-single cell RNA-seq analysis reveals the effects of anti-PD-L1 antibody on in vivo dynamics of tumour-reactive T cells.**

(a) Definition of ψ-Tocky Locus.
(b) Violin plots showing the expression of each of selected key differentially expressed genes (DEGs) in CD8 T single cells specifically increased in each ψ-Tocky Locus.
(c) Violin plots showing the expression of each of selected key DEGs in Tconv specifically increased in each ψ-Tocky Locus.
(d) Violin plots showing the expression of each of selected key DEGs in Treg in specifically increased in each ψ-Tocky Locus.
(e) Pathway analysis for CD8 T cells in each ψ-Tocky Locus.

**Extended Data 5: Blockade of PD-L1 and CTLA4 alters in vivo activities of tumour-infiltrating T cells following antigen recognition.**
Nr4a3-Tocky mice were treated with PD-L1 blocking antibody (aPD-L1), aCTLA-4 blocking antibody (aCTLA-4), the combination of aPD-L1 and aCTLA-4 (Combination), or Isotype Controls (IgG). Tumour-infiltrating T cells were analysed.

(a) Schematic figure outlining the experimental design of the study.
(b) Representative flow cytometry plots showing Timer Blue fluorescence (Blue, immature) vs Timer Red fluorescence (Red, mature) in Treg, Tconv, and CD8+ T cells from the three treatment groups. Thresholds were set using a fully stained WT mouse.

(c - e) Bar plot showing the percentage of Timer+ cells in (c) Treg, (d) Tconv, and (e)CD8+ T cells.

(f - h) Bar plots showing mean Angle values in (f) Treg, (g) Tconv, and (h)CD8+ T cells.

(i) Density plot showing the distribution of Treg across Timer Angle. Representative data of three independent experiments of n=3 each.
(j) Bar plots showing the frequency of Treg, Tconv, and CD8+ T cells in different TockyLoci.

Data were collected over 3 independent experiments with a total of n = 9 per group. Two-way ANOVA with Tukey's multiple comparisons test was applied. Dots represent individual data points, error bars represent +/- standard error.

Extended Data 6: **IC therapy reduces tumour-reactive eTreg and activates CD25high Tconv and CD8 T cell populations**

(a) Representation of the clusters identified by k-means on the PCA result of parameters selected for UMAP in the UMAP space Tumour infiltrating Treg.
(b) Representative flow cytometry plots showing *CD25* vs *OX40* gating of CD25hi OX40 hi 'eTreg' in tumour infiltrating Treg, and *Nr4a3*-Blue vs *Nr4a3*-Red of tumour-

infiltrating CD25hi OX40 hi 'eTreg' in mice treated with either IgG, aPD-L1, aCTLA4 and the combination of aPDL1+aCTLA4.

(c) Barplot representing the frequency of CD25hi OX40 hi 'eTreg' in tumour-infiltrating Treg (CD4+ Foxp3+) of mice treated with PDL-1 blocking antibody (aPDL1), aCTLA4 blocking antibody (aCTLA4), the combination of aPDL1 and aCTLA4 (Combination) and Isotype Controls (IgG).

(d) Barplots representing the frequency of Nr4a3 tocky+ cells in tumour-infiltrating CD25hi OX40 hi 'eTreg' and their mean Angle values in mice treated with PDL-1 blocking antibody (aPDL1), aCTLA4 blocking antibody (aCTLA4), the combination of aPDL1 and aCTLA4 (Combination) and Isotype Controls (IgG).

(e) Density plot representing *Nr4a3*-angle distribution in tumour-infiltrating CD25hi OX40 hi 'eTreg' of mice treated with PDL-1 blocking antibody (aPDL1), aCTLA4 blocking antibody (aCTLA4), the combination of aPDL1 and aCTLA4 (Combination) and Isotype Controls (IgG). Representative data of three independent experiments of n=3 each.

(f) Barplots representing the frequency of CD25hi OX40 hi 'eTreg' cells in different TockyLoci out of Timer+ CD25hi OX40 hi 'eTreg' cells of mice treated with PDL-1 blocking antibody (aPDL1), aCTLA4 blocking antibody (aCTLA4), the combination of aPDL1 and aCTLA4 (Combination) and Isotype Controls (IgG). Data was collected over 3 independent experiments with a total of n=9 per group.

(g) Representation of the clusters identified by k-means on the PCA result of parameters selected for UMAP in the UMAP space Tumour infiltrating Tconv.

(h) Representative flow cytometry plots showing *CD25* vs *OX40* gating of CD25hi OX40 hi Tconv in tumour infiltrating Tconv, and *Nr4a3*-Blue vs *Nr4a3*-Red of tumour-infiltrating CD25hi OX40 hi Tconv in mice treated with either IgG, aPD-L1, aCTLA4 and the combination of aPDL1+aCTLA4.

(i) Density plot representing *Nr4a3*-angle distribution in tumour-infiltrating CD25hi OX40 hi Tconv of mice treated with PDL-1 blocking antibody (aPDL1), aCTLA4 blocking antibody (aCTLA4), the combination of aPDL1 and aCTLA4 (Combination) and Isotype Controls (IgG). Representative data of three independent experiments of n=3 each.

(j) Barplots representing the frequency of CD25hi OX40 hi Tconv cells in different TockyLoci out of Timer+ CD25hi OX40 hi Tconv cells of mice treated with PDL-1 blocking antibody (aPDL1), aCTLA4 blocking antibody (aCTLA4), the combination of aPDL1 and aCTLA4 (Combination) and Isotype Controls (IgG). Data was collected over 3 independent experiments with a total of n=9 per group.

(k) Representation of the clusters identified by k-means on the PCA result of parameters selected for UMAP in the UMAP space Tumour infiltrating CD8+ T.

(l) Representative flow cytometry plots showing *CD25* vs *PD1* gating of CD25hi PD1hi CD8+ T in tumour infiltrating CD8+ T cells, and *Nr4a3*-Blue vs *Nr4a3*-Red of tumour-infiltrating CD25hi PD1hi CD8+ T cells in mice treated with either IgG, aPD-L1, aCTLA4 and the combination of aPDL1+aCTLA4.

(m) Density plot representing *Nr4a3*-angle distribution in tumour-infiltrating CD25hi PD1hi CD8+ T cells of mice treated with PDL-1 blocking antibody (aPDL1), aCTLA4 blocking antibody (aCTLA4), the combination of aPDL1 and aCTLA4 (Combination) and Isotype Controls (IgG). Representative data of three independent experiments of n=3 each.

(n) Barplots representing the frequency of CD25hi PD1hi CD8+ T cells in different TockyLoci out of Timer+ CD25hi PD1hi CD8+ T cells of mice treated with PDL-1 blocking antibody (aPDL1), aCTLA4 blocking antibody (aCTLA4), the combination of aPDL1 and aCTLA4 (Combination) and Isotype Controls (IgG). Data was collected over 3 independent experiments with a total of n=9 per group.

Extended Data 7: **Addition of an OX40 agonist to the combination further increases effector anti tumour T cell populations and decreases anti-tumour regulatory populations**

(a) Density plot representing *Nr4a3*-angle distribution in tdLN Treg cells of mice treated with the double combination of aPDL1 and aCTLA4, the triple combination of aPDL1, aCTLA4 and aOX40 and Isotype Controls (IgG) (n=6 each).

(b) Barplots representing the frequency of tdLN Treg cells in different TockyLoci out of Timer+ Treg of mice treated with the double combination of aPDL1 and aCTLA4, the triple combination of aPDL1, aCTLA4 and aOX40 and Isotype Controls (IgG).

(c) Representation of the clusters identified by k-means on the PCA result of parameters selected for UMAP in the UMAP space tdLN Treg.

(d) Violin plots representing the Intensity mean of clusters generated by cluster fitting of tdLN Tconv.

(e) Representative flow cytometry plots of the gating strategy to identify anti-tumour Treg populations across treatments, this includes PD1+ CD62L- GITRhi ICOShi Treg cells and their Nr4a3-Blue vs Red pattern.

(f) Barplot representing the frequency of PD1+ CD62L- Treg out of the tdLN Treg population of mice treated with the double combination of aPDL1 and aCTLA4, the triple combination of aPDL1, aCTLA4 and aOX40 and Isotype Controls (IgG).

(g) Representation of the clusters identified by k-means on the PCA result of parameters selected for UMAP in the UMAP space tdLN Tconv.

(h) Violin plots representing the Angle mean of clusters generated by cluster fitting of tdLN CD8+ T.

(i) Representative flow cytometry plots of the gating strategy to identify anti-tumour Tconv populations across treatments, this includes PD1+ CD62L- ICOShi CXCR5hi GITRhi Tconv cells and their Nr4a3-Blue vs Red pattern.

(j) Barplot representing the frequency of PD1+ CD62L- Tconv out of the tdLN Tconv population of mice treated with the double combination of aPDL1 and aCTLA4, the triple combination of aPDL1, aCTLA4 and aOX40 and Isotype Controls (IgG).
(k) Representation of the clusters identified by k-means on the PCA result of parameters selected for UMAP in the UMAP space tdLN CD8+ T.
(l) Violin plots representing the Intensity mean of clusters generated by cluster fitting of tdLN CD8+ T.
(m) Representative flow cytometry plots of the gating strategy to identify anti-tumour CD8+ T populations across treatments, this includes PD1+ CD62L- ICOShi CD69hi CD8+ T cells and their Nr4a3-Blue vs Red pattern.
(n) Barplot representing the frequency of PD1+ CD62L- CD8+ T out of the tdLN CD8+ T population of mice treated with the double combination of aPDL1 and aCTLA4, the triple combination of aPDL1, aCTLA4 and aOX40 and Isotype Controls (IgG).

**Extended Data 8: The OX40 agonist enhances the effects of the combination checkpoint blockade on the tumour-infiltrating, reactive T cells in a unique manner**

(a) Representation of the clusters identified by k-means on the PCA result of parameters selected for UMAP in the UMAP space of tumour-infiltrating Treg.
(b) Violin plots representing the Angle and Intensity means of clusters generated by cluster fitting of Tumour-infiltrating Treg cells
(c) Representation of the clusters identified by k-means on the PCA result of parameters selected for UMAP in the UMAP space of tumour-infiltrating Tconv.
(d) Violin plots representing the Angle and Intensity means of clusters generated by cluster fitting of Tumour-infiltrating Tconv cells
(e) Representative flow cytometry plots of the gating strategy to identify UMAP cluster 3 Tconv population across treatments, this includes PD1hi GITRhi ICOShi CD69hi Tconv cells and their Nr4a3-Blue vs Red pattern.
(f) Representative flow cytometry plots of the gating strategy to identify UMAP cluster 2 Tconv population across treatments, this includes PD1lo KLRG1+Tconv cells and their Nr4a3-Blue vs Red pattern.

Extended Data 9: **The OX40 agonist enhances the effects of the combination checkpoint blockade on the tumour-infiltrating, reactive CD8 T cells in a unique manner**

(a) Representation of the clusters identified by k-means on the PCA result of parameters selected for UMAP in the UMAP space of tumour-infiltrating CD8 T cells.

(b) Representative flow cytometry plots of the gating strategy to identify UMAP cluster 4 CD8+ T population across treatments, this includes PD1hi GITRhi CD69hi CD25hi KLRG1- CD62L- cells and their Nr4a3-Blue vs Red pattern.

(c) Representative flow cytometry plots of the gating strategy to identify UMAP cluster 8 CD8+ T population across treatments, this includes PD1hi CD62L- KLRG1+ cells and their Nr4a3-Blue vs Red pattern.
