## Supplementary figures and images for "Single-cell level temporal profiling of tumour-reactive T cells under immune checkpoint blockade"

### Extended Data 1

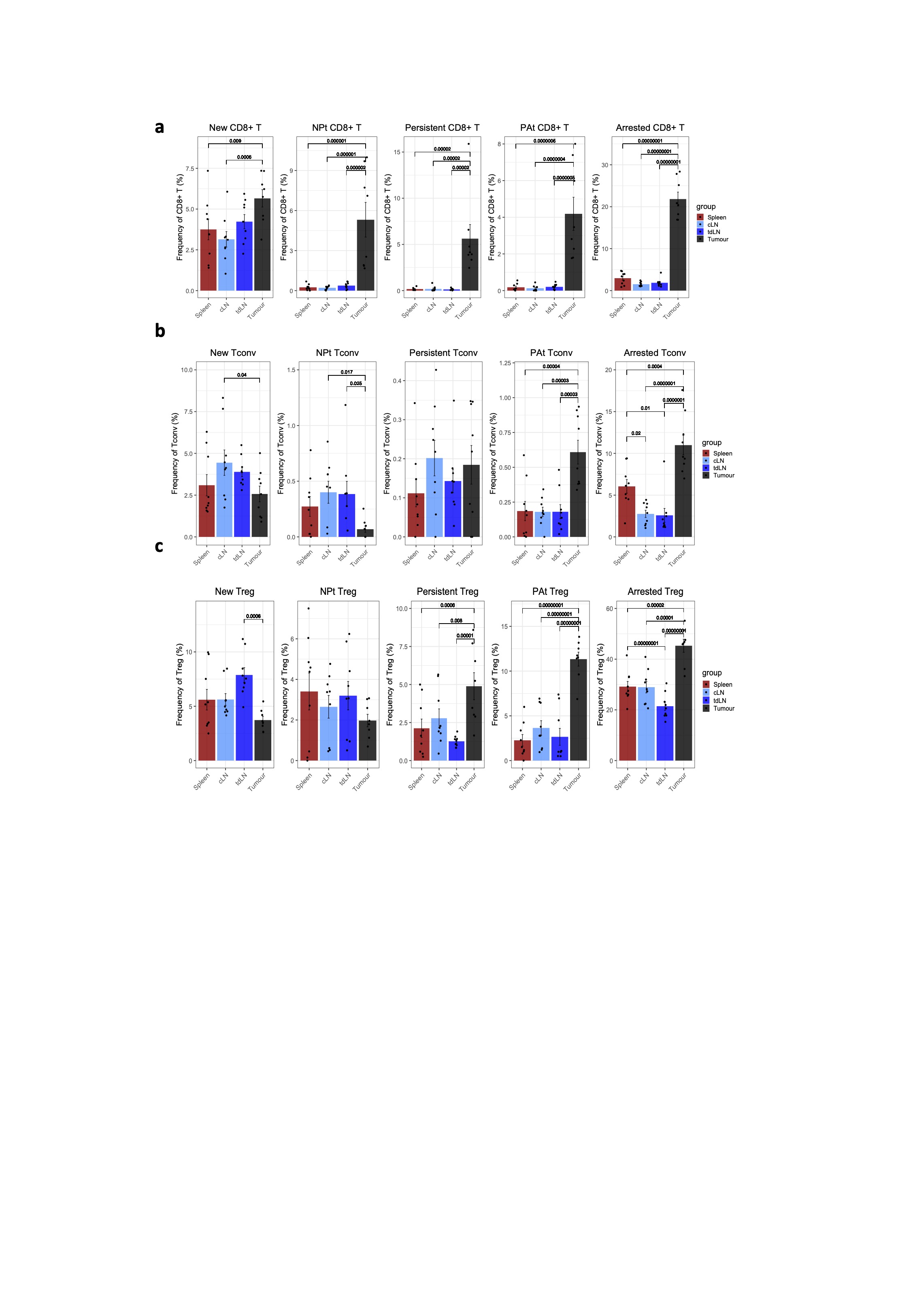

### Extended Data 2

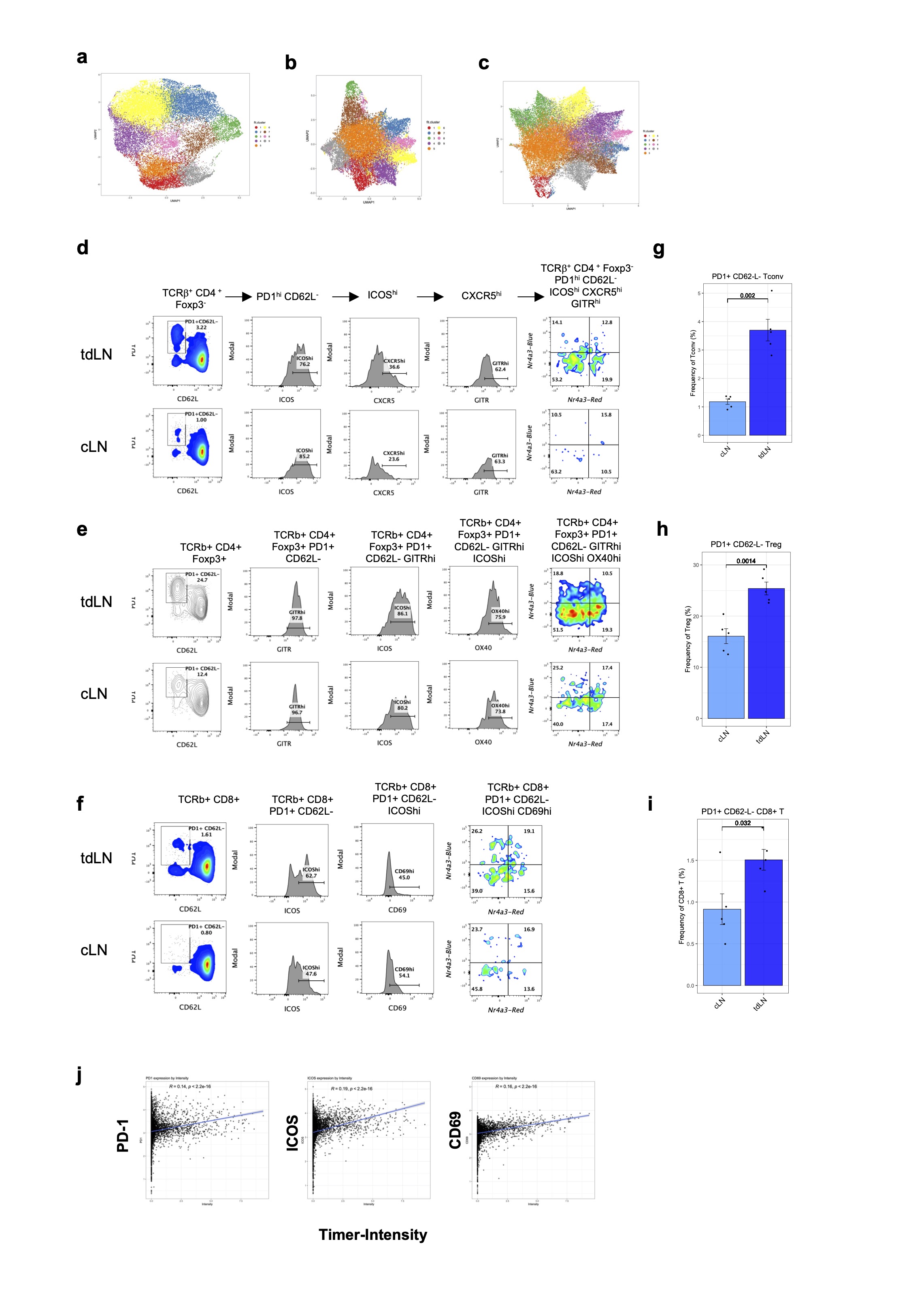

### Extended Data 3

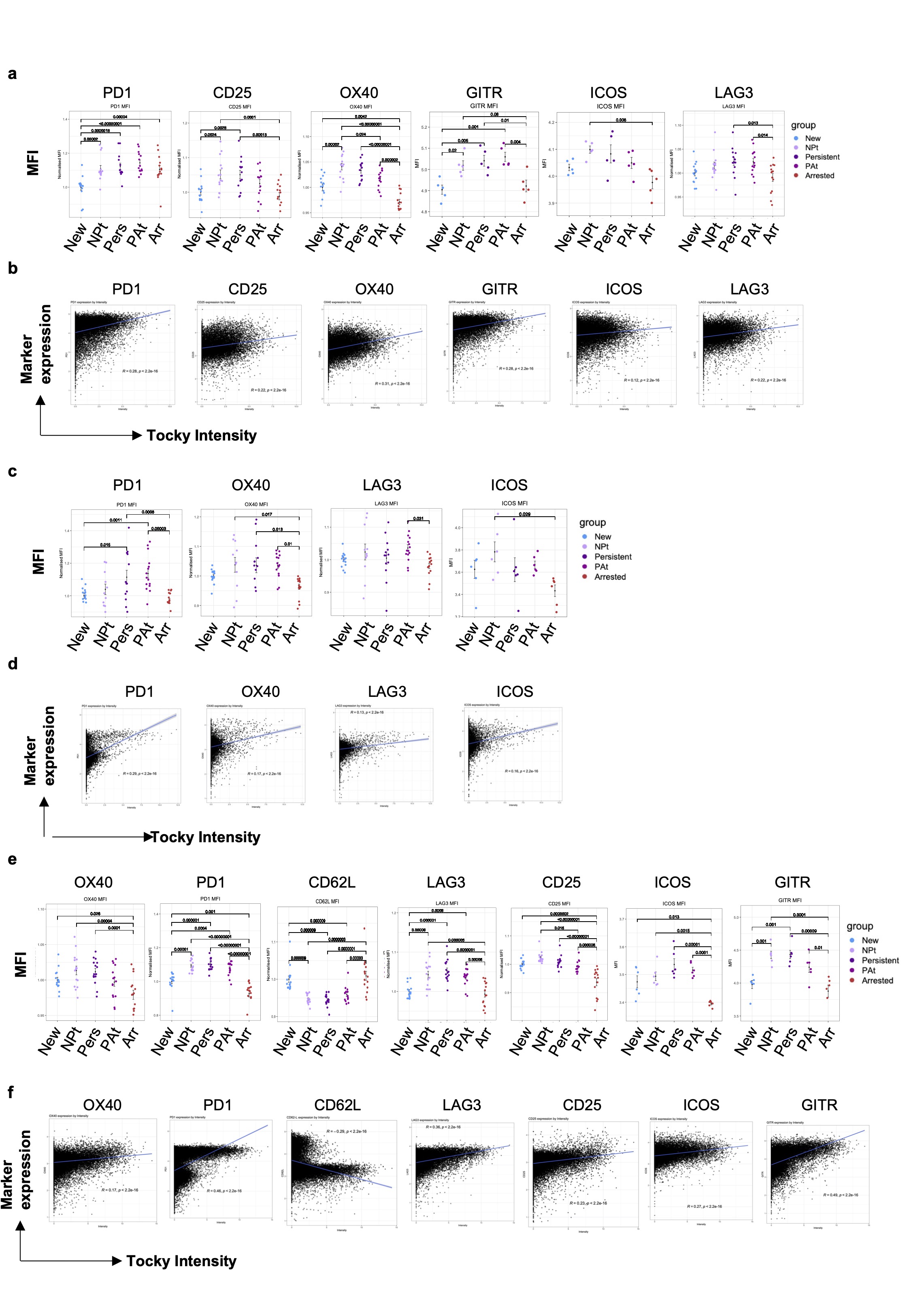

### Extended Data 4

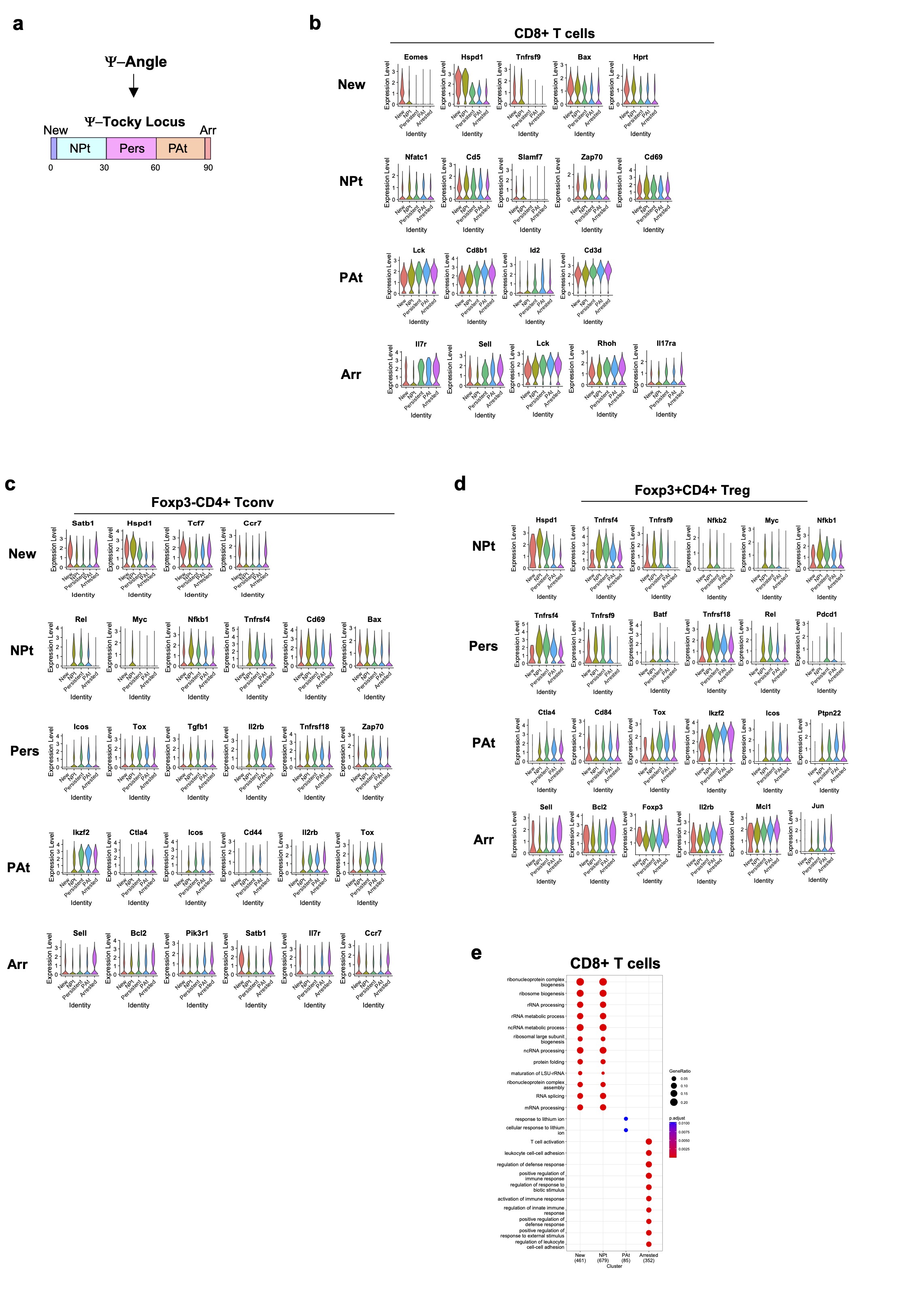

### Extended Data 5

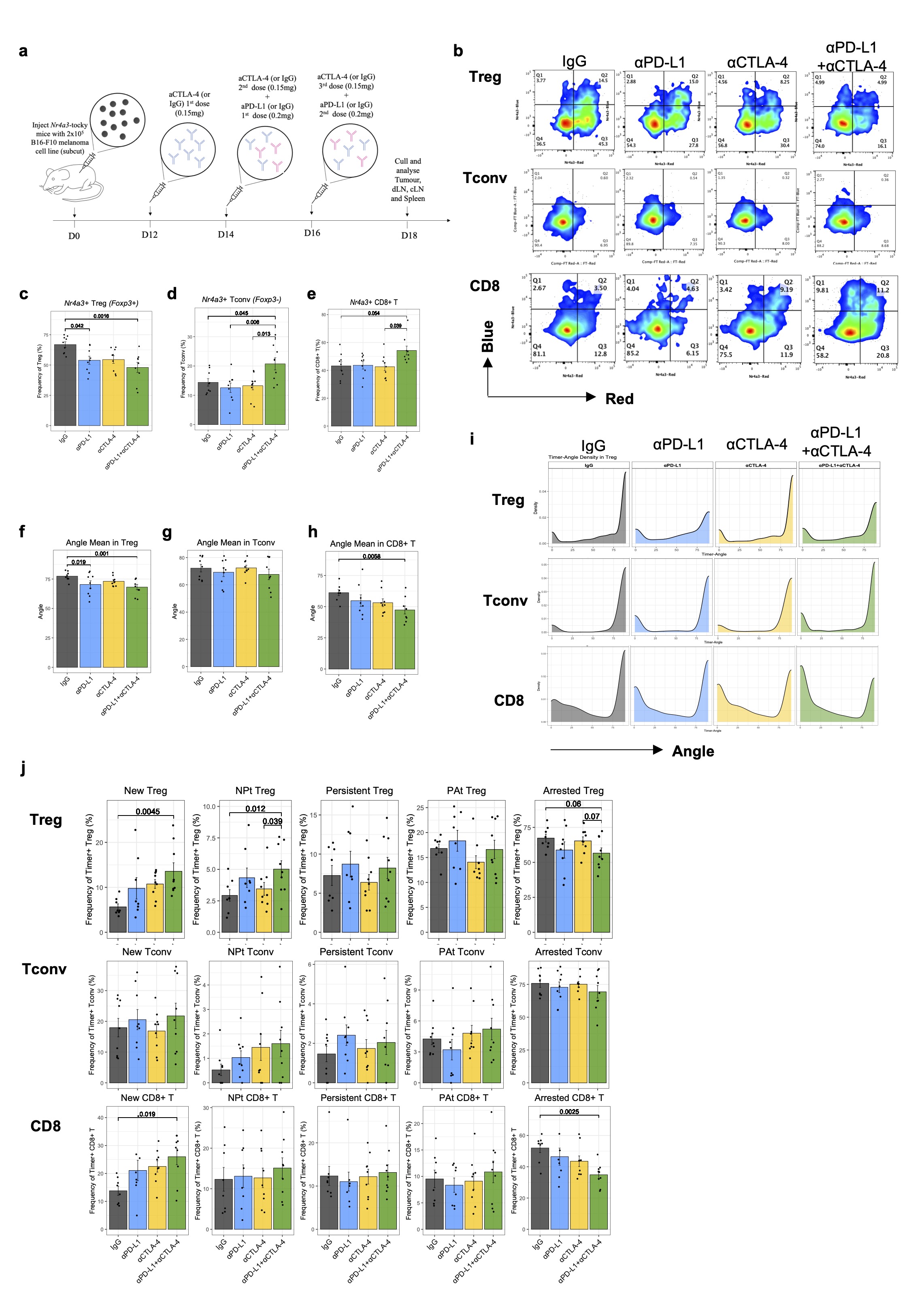

### Extended Data 6

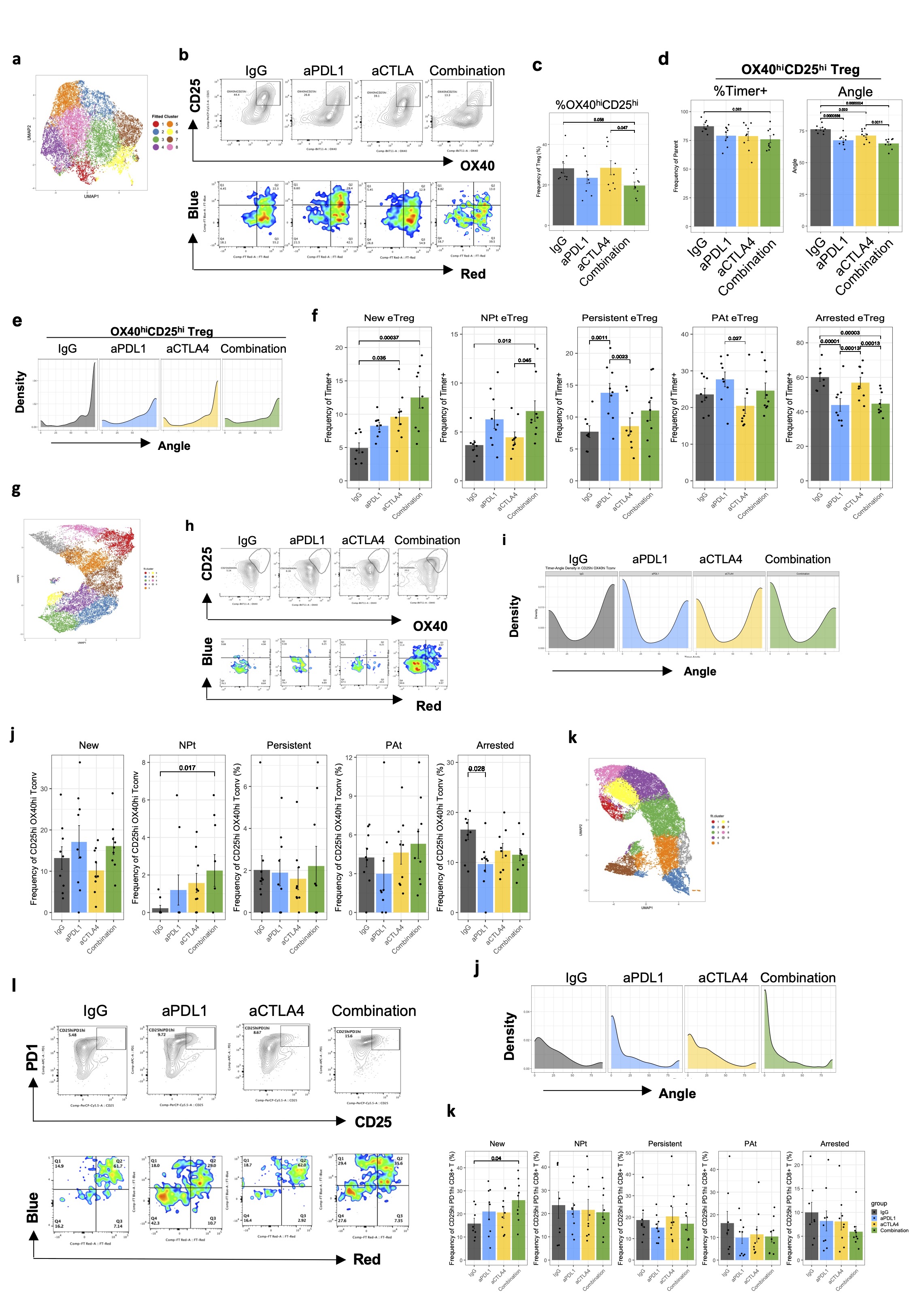

### Extended Data 7

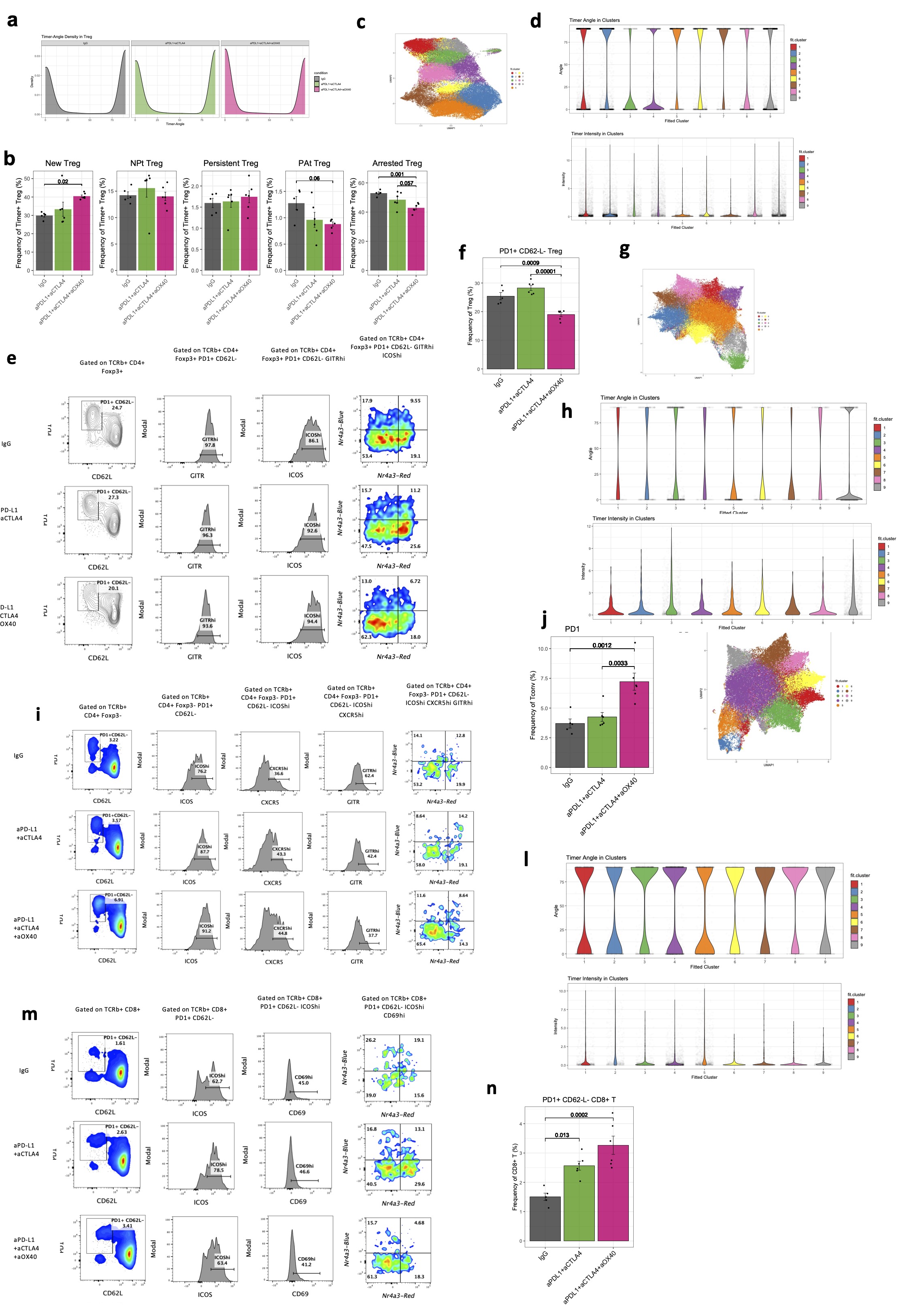

### Extended Data 8

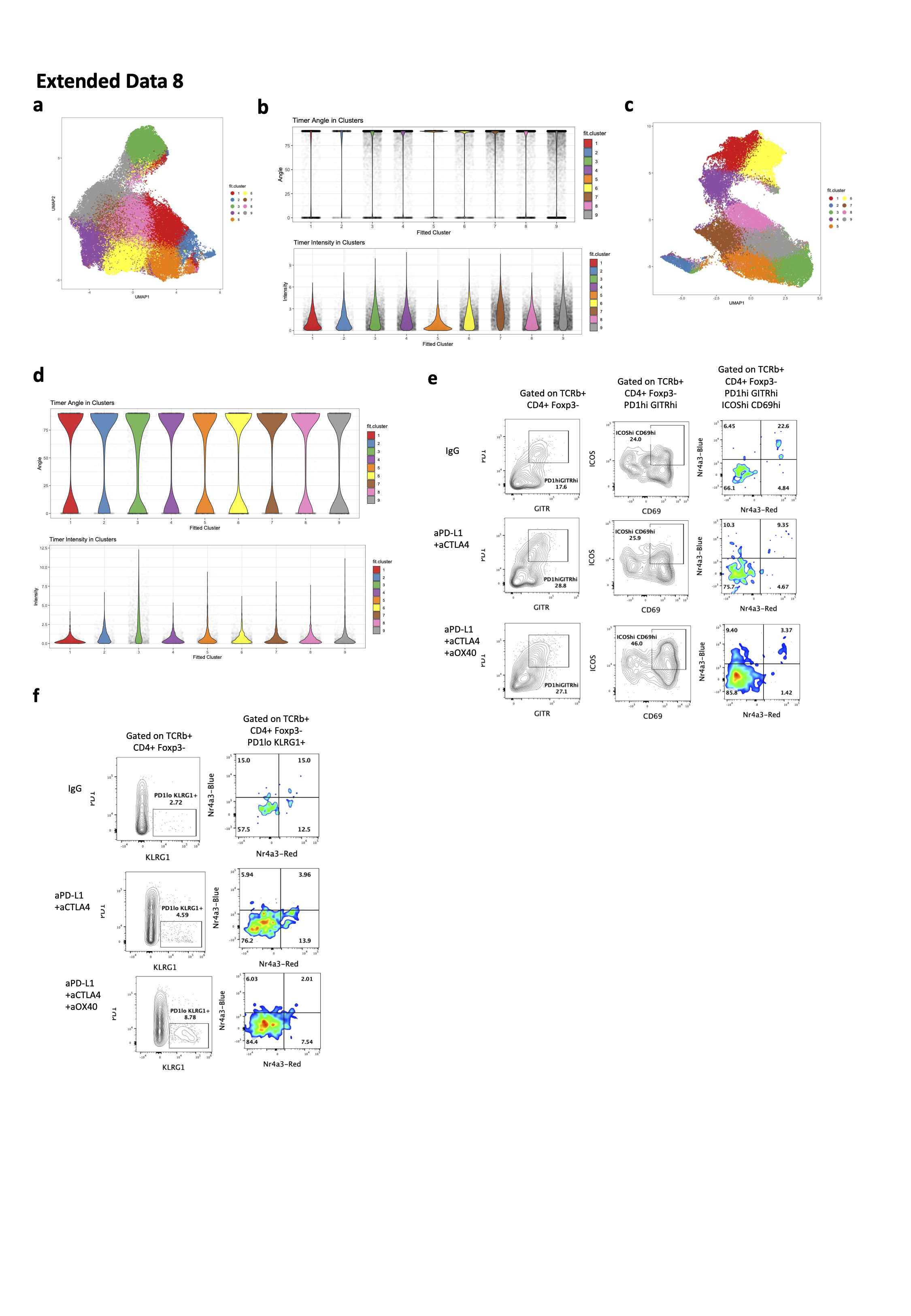

### Extended Data 9

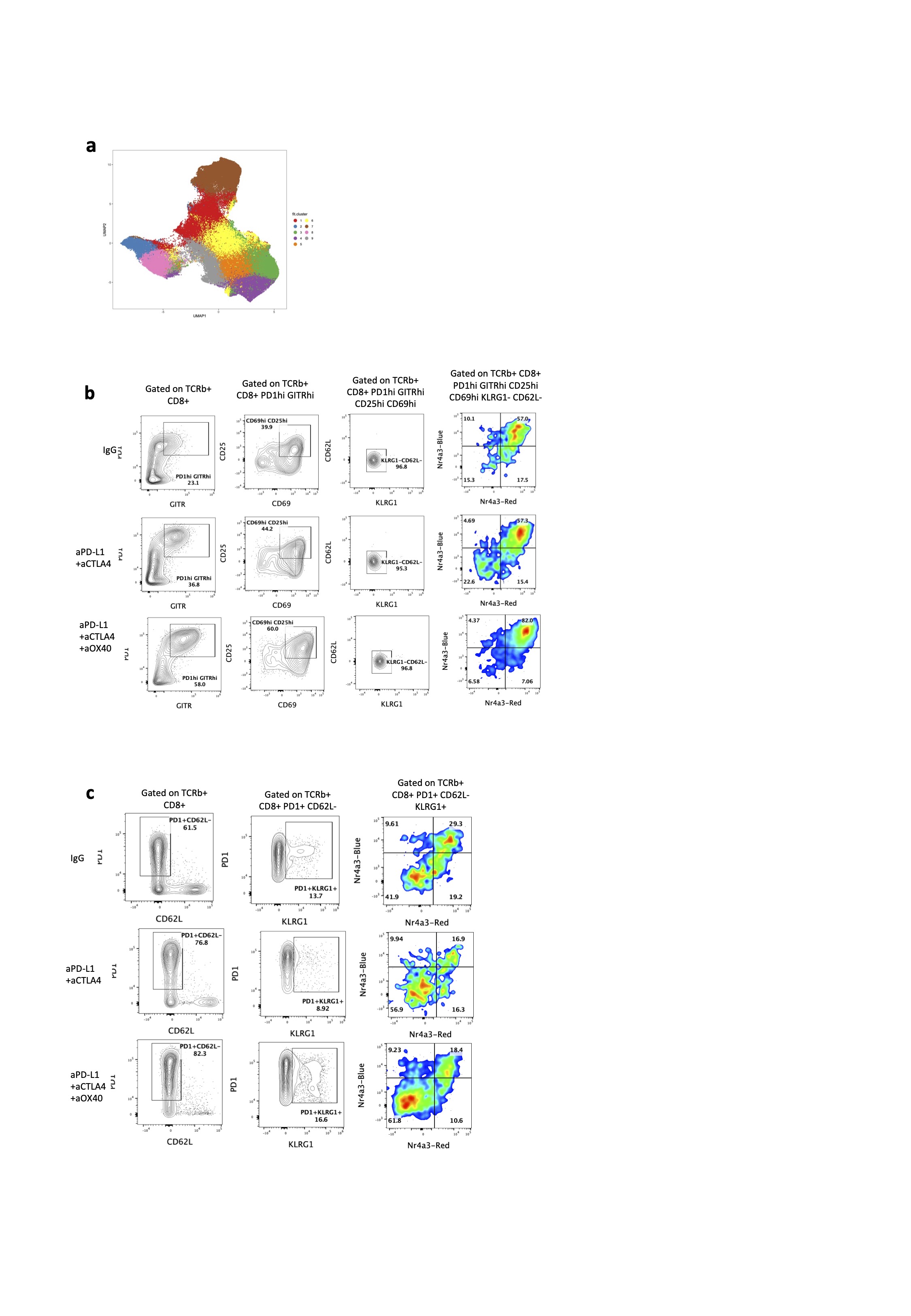
